# Electroencephalography, pupillometry, and behavioral evidence for locus coeruleus-noradrenaline system related tonic hyperactivity in older adults

**DOI:** 10.1101/2025.10.02.680040

**Authors:** Andy Jeesu Kim, Santiago Morales, Joshua Senior, Mara Mather

**Author notes:** Corresponding author:* Andy Jeesu Kim, 3715 McClintock Ave. Los Angeles, CA 90089.

## Abstract

Neuroimaging studies have shown that age-related dysregulation of the locus coeruleus-noradrenaline (LC-NA) system is associated with cognitive decline. However, due to limitations in directly measuring LC function *in vivo*, it remains unclear whether age-related alterations in humans reflect tonic LC-NA system hyper- or hypoactivity, constraining our understanding of underlying mechanisms and hampers the development of targeted preventative interventions. In this study, we tested the hypothesis that cognitively healthy older adults sustain tonic LC hyperactivity, by acquiring electrophysiological, pupillometric, and behavioral measures during a passive and active auditory oddball paradigm. We capitalized on the LC-NA system’s role in arousal regulation and manipulated state arousal using the unpredictable threat of electric shock. We hypothesized that if older adults maintain elevated LC activity compared with young adults, task-evoked noradrenergic responses would be less responsive to arousal in older adults. Consistent with this hypothesis, arousal elicited weaker behavioral responses, pupil dilation responses, and P300 event-related potentials in older adults compared with young adults. Linear mixed models revealed an arousal by modality interaction, showing that arousal differentially modulated attentional control to salient but task-irrelevant distractors between both age groups. Collectively, these findings support the hypothesis that aging is associated with tonic LC-NA system hyperactivity in humans, with neuromodulatory consequences for mechanisms of attentional control. Furthermore, the multimodal approach underscores the potential of non-invasive physiological markers to assess LC-NA system function throughout aging and identify individuals at elevated risk for neurodegenerative progression prior to the emergence of clinical biomarkers.

**Highlights:**

- Behavioral, pupil and EEG data indicate LC-NA tonic hyperactivity in older adults.
- Arousal modulates attentional control differently in young vs. older adults.
- Older adults show increased distractibility from reduced habituation.
- Arousal differentially modulates alpha power and aperiodic activity across age groups.
- Phasic noradrenergic responses are modulated by state, arousal, and attention.

## Introduction

Normal and pathological aging are associated with declines across multiple cognitive domains, including memory, language, executive function, processing speed, and visuospatial abilities (Bondi et al., 2014; Buckner, 2004; Jak et al., 2009; Murman, 2015). These cognitive changes often coincide with declines in both gray and white matter, as well as functional changes that are frequently interpreted as compensatory mechanisms (Fjell & Walhovd, 2010; Bishop et al., 2010; Damoiseaux, 2017). In addition, disruptions in neuromodulatory systems have also shown to be implicated in age-related cognitive decline, with evidence of altered structural and functional connectivity between neuromodulatory nuclei (Krohn et al., 2023; Slater et al., 2022; Weinshenker, 2018), and significant neuronal loss (Lyness et al., 2003; Orlando et al., 2023). Specifically, the locus coeruleus-noradrenaline (LC-NA) system, which supports a broad range of cognitive functions (Aston-Jones et al., 1999; Aston-Jones & Cohen, 2005; Bouret & Sara, 2005), plays a critical role in maintaining cognitive health across the lifespan (Clewett et al., 2016; Mather & Harley, 2016; Wilson et al., 2013). However, the LC is the earliest region to exhibit neuropathology as observed in postmortem brains (Braak et al., 2011) and tau accumulation in the medial temporal lobe is preceded by declines in LC MRI structural contrast (Bueichekú et al., 2024), suggesting neuropathological spread from the LC. The accumulation of LC neurodegenerative pathology is associated with cognitive impairment (Grudzien et al., 2007; Matthews et al., 2002; Nelson et al., 2012; Theofilas et al., 2017). Noninvasive magnetic resonance imaging (MRI) structural sequences produce high contrast at the location of noradrenergic LC neurons (Keren et al., 2015; Watanabe et al., 2019) and so provide a way to assess LC structure *in vivo*. In older adults, reduced LC MRI structural contrast has been linked to cognitive decline and increased risk for neurodegenerative diseases (Cassidy et al., 2022; Clewett et al., 2016; Dahl et al., 2023; Dahl, Mather, et al., 2019; Dahl, Mather, Werkle-Bergner, et al., 2022; Elman et al., 2021; Hämmerer et al., 2018; Liu et al., 2020). Neuroimaging studies have identified that LC MRI structural contrast increases in early adulthood, peaks in midlife, and subsequently declines in late life (Liu et al., 2019). Furthermore, recent studies have identified that LC MRI structural contrast in older adults predicts tau spread in subsequent years (Bueichekú et al., 2024), and also predicts future memory performance (Dahl et al., 2023). Thus, being able to measure age-related changes in the LC-NA system function in cognitively healthy older adults may provide a critical window for detection and preventative interventions aimed at mitigating cognitive decline prior to significant development of neuropathology.

One currently open question is whether LC tonic activity increases or decreases with age. Initial findings with rats were mixed, showing significantly lower tonic activity (Olpe & Steinmann, 1982) or non-significantly higher tonic activity (Shirokawa et al., 2000) in older rats than younger rats. More recently, over 1000 recordings pooled across labs of rats’ single neurons indicate older rats show greater LC neuronal tonic activity than younger rats (Kelberman et al., 2024). In human AD patients, degeneration of LC neurons is accompanied by increased metabolic activity in surviving neurons (as indicated by an increased ratio of the noradrenaline metabolite 3-methoxy-4-hydroxyphenylglycol or MHPG to noradrenaline; Hoogendijk et al., 1999), suggesting that noradrenergic hyperactivity may develop to compensate for neuronal loss (Liu et al., 2025; Szot et al., 2006). In contrast, in the TgF344-AD rat model of early-onset AD, tonic LC activity is lower at 6 and 15 months than in wild-type rats (Kelberman et al., 2023). Because in the TgF344-AD rat, LC exhibits hyperphosphorylated tau before the entorhinal cortex or hippocampus, it has been argued that this rat model may provide a model of human AD, in which hyperphosphorylated (pretangle) tau in the LC is the first detected sign of AD pathology in the brain. However, a significant difference between this rat model and the human AD cases is that, in the rat model, the hyperphosphorylated tau is a response to a genetic mutation affecting Aβ concentrations (Cohen et al., 2013) and so far has only been detected at a point when Aβ plaques are already present in cortex (Rorabaugh et al., 2017), whereas in humans, LC hyperphosphorylated tau typically emerges decades before Aβ plaque. These divergent patterns underscore the further need to investigate changes in LC function in humans as the trajectory of LC dysregulation appears to differ between rodent models of AD and human AD.

However, direct measurement of LC activity in humans still poses a significant challenge (Kim, 2023). The field currently lacks a reliable, absolute measure of LC function in humans and has therefore largely relied on structural intensity contrasts to investigate changes in LC structure (Betts et al., 2019; Liu et al., 2020). More recent studies have employed ultra-high-field neuroimaging at 7 Tesla to improve signal-to-noise in the brainstem, but these investigations are scarce due to limited accessibility (Berger et al., 2023; Boukezzi et al., 2025; Ludwig et al., 2024). Finally, Bang et al. (2023) have measured sub-second noradrenergic dynamics using clinical depth electrodes implanted in the amygdala of epilepsy patients and used an electrochemical model approach to link noradrenergic activity to arousal and attention in a visual affective oddball task (Bang et al., 2023). With only invasive, expensive, and unique circumstances allowing for its direct measurement in humans, the trajectory of noradrenergic function in human aging remains poorly understood and it is yet unclear whether humans exhibit LC-NA system hyperactivity during aging to the presence of neuropathology in the LC (Braak et al., 2011).

Therefore, in this study, we examined age-related changes in LC-NA system activity in cognitively healthy older adults using multiple modalities that are influenced by LC activity including behavior, pupillometry, and electroencephalography (EEG). Building on evidence from animal studies of LC activity in aging (Kelberman et al., 2024), we hypothesized that human aging is characterized by LC hyperactivity. Although direct measurement of LC activity in humans is methodologically challenging, pupillometry has been widely used as an indirect proxy of noradrenergic function (Costa & Rudebeck, 2016; Elman et al., 2017; Gilzenrat et al., 2010; Joshi & Gold, 2020; Joshi et al., 2016; Kremen et al., 2019; Liu et al., 2017; Murphy et al., 2011; Weigand et al., 2023). Specifically, pupil dilation responses to stimuli have been directly linked to phasic or bursts of noradrenergic activity (Breton-Provencher & Sur, 2019; DiNuzzo et al., 2019; Grueschow et al., 2022, 2021; Joshi et al., 2016; Liu et al., 2017; Murphy et al., 2014), with robust phasic responses at optimal levels of tonic, or baseline, LC firing (Aston-Jones & Cohen, 2005; Hayat et al., 2020). In addition, the P300 component of the event-related potential (ERP) has been associated with phasic noradrenergic activity (Foote et al., 1991; Murphy et al., 2011; Nieuwenhuis et al., 2005; Swick et al., 1994), specifically in the three-stimulus oddball paradigm (Polich, 2007). To test our hypothesis, we simulated altered noradrenergic states and activity by employing the threat of unpredictable shock paradigm (Schmitz & Grillon, 2012) and leveraged the LC-NA system’s functional role in mediating state arousal (Breton-Provencher & Sur, 2019; Clewett et al., 2018; Mather & Harley, 2016; Murphy et al., 2011). Previous findings have shown that exposure to the threat of unpredictable shock has been found to suppress parietal alpha oscillations (Balderston et al., 2017), making young adults’ parietal alpha oscillations more closely resemble neural patterns typically observed in older adults (Vaden et al., 2012), suggesting that this model of arousal may emulate changes in LC dynamics that occur during aging. In rodents, LC tonic and phasic activity show a negative linear association, such that the strongest phasic responses occur during low tonic states (e.g., NREM sleep) and the weakest phasic responses are observed during high tonic states (e.g., active wakefulness; Hayat et al., 2020). Therefore, we predicted that young adults characterized by optimal baseline LC firing would show reduced performance under sustained threat due to overactivation of the arousal system (e.g., increased tonic LC activity). In contrast, we hypothesized that older adults, if characterized by elevated tonic noradrenergic hyperactivity, would show reduced sensitivity to arousal induction.

In this experiment, we collected resting-state EEG data to examine both oscillatory and aperiodic activity, as well as stimulus-evoked ERPs during a three-stimulus auditory oddball paradigm. EEG recordings were obtained in a neutral condition and under the threat of *unpredictable* electric shock (Schmitz & Grillon, 2012). To assess mechanisms of attentional control, participants completed both passive and active versions of the oddball task. In the passive condition, participants were instructed to ignore the tones, whereas in the active condition, they were required to respond to all tones with a button press but used a distinct response only for target tones (Mertens & Polich, 1997; Wronka et al., 2008). In addition to EEG, we measured baseline pupil diameter during resting-state recordings and stimulus-evoked pupil responses to standard, target, and distractor tones during the task. Previous research has shown that the P300 ERP component is closely related to pupil dilation responses (Murphy et al., 2011; Nieuwenhuis et al., 2005; Podvalny et al., 2021), and that alpha oscillations are functionally linked to P300 amplitude (Studenova et al., 2023; Yordanova & Kolev, 1998). Aging has been reliably associated with reduced P300 amplitude and delayed peak latencies (O’Connell et al., 2012; Polich, 1997). Moreover, age-related EEG changes include reduced aperiodic-adjusted alpha power and a slowing of peak alpha frequency (Donoghue et al., 2020a; Merkin et al., 2023; Park et al., 2025; Tröndle et al., 2023), with higher resting-state alpha power linked to improved spatial memory performance (Jabès et al., 2021). Aperiodic signals are characterized by measures of an exponent (1/f slope of power spectrum) which reflects the balance of excitatory and inhibitory cortical activity from AMPA and GABA receptor-mediated currents, and aperiodic offset which reflects shifts in the broadband power across all frequencies (Donoghue et al., 2020; Gao et al., 2017). Importantly, these signals have been associated with both neuronal population spiking and fMRI BOLD signals (Donoghue et al., 2020; Gao et al., 2017; Jacob et al., 2021; Park et al., 2025), demonstrating that these non-oscillatory signals reflect meaningful physiological processes rather than signatures of noise as previously considered. Interestingly, while aperiodic activity changes with healthy aging, it appears to remain stable during the progression of AD, when only oscillatory power continues to decline (Kopčanová et al., 2024). Based on our hypothesis, we predicted exposure to the threat of shock would suppress alpha power and alter the aperiodic exponent in young adults, mimicking neural signatures commonly associated with aging (Balderston et al., 2017; Donoghue et al., 2020). We further expected that this arousal manipulation would reduce P300 amplitudes and delay peak latencies as seen in young adults (Chang et al., 2024; MacNamara & Barley, 2018). However, we hypothesized that cognitively healthy older adults would show relatively smaller changes compared with young adults, consistent with the notion of sustained noradrenergic hyperactivity in aging.

## Methods

### Participants

Data from older adults were obtained from a randomized controlled trial (ClinicalTrials.gov Identifier: NCT05602220) examining the effects of heart rate and breathing regulation on attention and memory (Nashiro et al., 2024). Specifically, we used baseline data collected during Visit 1, prior to the initiation of the intervention. Older adults were recruited from the local Los Angeles communities for monetary compensation. In the present study, we included all data from 66 older adults (Age: mean = 60.2 years old, SE = 0.7; Gender: 36 female, 30 male; Ethnicity: Asian – 12, Bi-racial – 6, Black – 8; Undisclosed - 4, White - 36) who completed the EEG recording session. To examine age-related differences, we additionally recruited a comparison group of young adults through the University of Southern California (USC) SONA subject pool for course credit. The young adult sample consisted of 68 participants (Age: mean = 19.8 years old, SE = 0.2; Gender: 41 female, 25 male, 2 undisclosed; Ethnicity: American Indian – 1, Asian – 30, Bi-Racial – 2, Black - 3, Other – 7, Undisclosed – 6 White – 19).

### Apparatus

Stimuli were presented from a custom-built NZXT desktop computer (NZXT, Los Angeles, CA, USA) running MATLAB 2024b (Mathworks, Natick, MA, USA) with Psychophysics Toolbox extensions (Brainard, 1997). Visual stimuli were displayed on a Sun Microsystems 4472 CRT monitor (Oracle Corporation, Santa Clara, CA, USA) with a refresh rate of 85 Hz. Pupillometry data were acquired using the EyeLink 1000 Plus system (SR Research Ltd., Ottawa, Ontario, Canada) and participants’ head position were stabilized using a chin rest from SR Research. Aversive electrical stimulation was delivered via a battery-powered transcutaneous stimulator (Model E13-22; Coulbourn Instruments, Allentown, PA, USA) to the third and fourth fingers. Electrodes (Model EL507A; BIOPAC Systems, Goleta, CA, USA) were attached using conductive electrode gel (GEL101A; BIOPAC Systems). Auditory stimuli were presented through insert earphones designed for research (Model ER2; Etymotic Research, Inc., Fort Worth, TX, USA) with disposable foam ear tips (Models ER3-14B and ER3-14C; Etymotic Research).

All experimental procedures took place in a soundproof and electromagnetically shielded testing room constructed as a Faraday cage. Electrical equipment was powered externally and communication between the participant and experimenter was maintained via a two-way night-vision baby monitor system (Model SM935A; Kidsneed). EEG data were recorded using a 65-channel HydroCel Geodesic Sensor Net, sampled at 1000 Hz, and acquired via Net Station software (Version 5.4; Electrical Geodesics, Inc., Eugene, OR, USA). Channel Cz served as the online reference during data collection and data were re-referenced offline to the average of the mastoid electrodes during data pre-processing. Electrode impedance was continuously monitored throughout the session and maintained below 50 kΩ.

### Design and Procedure

The experiment procedure was conducted under both no-threat (i.e., control) and threat of unpredictable shock conditions (see Figure 1). Each block began with a ten-minute resting-state EEG recording, consisting of two five-minute blocks of eyes closed and open, respectively. During the eyes open condition, participants were instructed to fixate on a central fixation cross (0.7° x 0.7° visual angle). Following resting-state recordings, participants completed two runs of the passive oddball task. Participants were instructed to disregard the auditory stimuli and look at the central fixation cross. Each run consisted of 120 trials composed of 70% standard tones (500 Hz), 15% target tones (1000 Hz), and 15% distractor tones (salient auditory tone; burst of white noise), all presented at 75 dB. Inter-stimulus intervals (ISI) were jittered at 2.2, 2.3, or 2.4 seconds, and randomly distributed across trials equally often. Following the passive task, participants underwent a calibration procedure to determine the subjective threshold for electric shock. Shocks were calibrated to an intensity deemed “unpleasant, but not painful” using a stepwise procedure beginning from the lowest setting and gradually increasing until the participant verbally confirmed the intensity threshold as in prior studies (Kim & Anderson, 2020a, 2020b). Participants were only connected to the shock device during this calibration process or when completing the shock block to allow for the arousal-inducing nature of the device to dissipate (Kim & Anderson, 2020a). Participants then repeated the resting-state recording and passive oddball task under the threat of shock condition. During the resting-state recordings, a single electric shock was randomly delivered every minute. During the passive oddball task, shocks were randomly administered twice within every set of 20 trials. Importantly, shock delivery took the place of a trial, so it never interfered with task performance. After the passive task block under threat of shock, participants were disconnected from the shock device. Finally, participants practiced the active oddball task in which they were instructed to press the bottom right button on the button box (SR Research) upon hearing the target tone and the middle button for all other tones. Thus, participants were only required to pay attention to the target tone but still made a manual response to all presented tones. Participants were required to achieve at least 90% accuracy during the practice run to verify understanding of task instructions. The active oddball task was identical in structure and timing to the passive task, with the only difference being the task instructions. Participants then completed two runs of the active oddball task under no-threat conditions, followed by two additional runs under the threat of shock condition. The frequency and randomization scheme of shock delivery in the active task mirrored that of the passive task.

**Figure 1.**
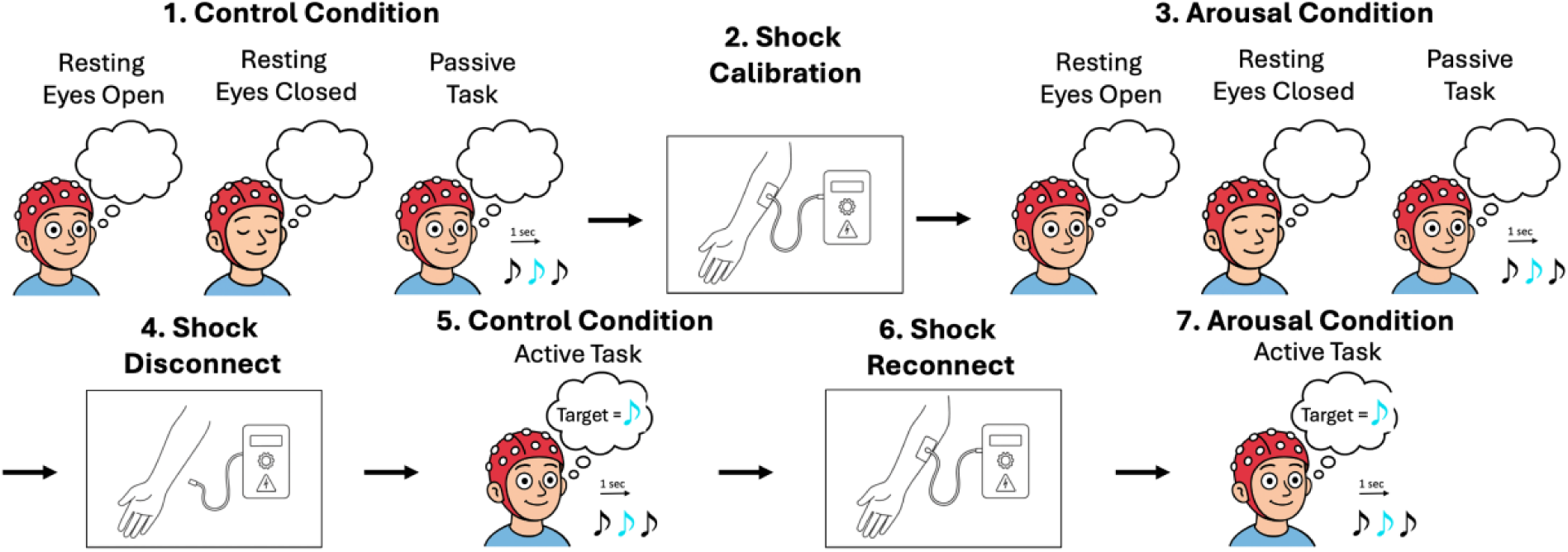
Experiment Task Sequence. EEG recordings were obtained from both young and older adults during resting-state conditions (eyes open and eyes closed, 5 minutes each) and while performing a three-stimulus auditory oddball task in both passive and active versions. All tasks were completed under a no-threat condition (control) and under the threat of unpredictable shock (arousal). The order of tasks was selected to reduce the influence of fatigue when comparing across arousal conditions.

### EEG Data Pre-processing

EEG data were processed using the standardized and semi-automated MADE pipeline (Debnath et al., 2020; Leach et al., 2020), as previously described (Morales et al., 2022). Raw continuous EEG data were first exported to MATLAB for offline preprocessing using the EEGLAB toolbox (Delorme & Makeig, 2004). Data were down sampled to 500 Hz, high-pass filtered at 0.1 Hz, and low-pass filtered at 50 Hz. Artifact-laden channels were identified and removed using the FASTER algorithm (Nolan et al., 2010). Next, independent component analysis (ICA) was performed by first creating a copy of the dataset. This copy was high-pass filtered at 1 Hz and segmented into 1-second epochs. Noisy segments and EMG artifacts were excluded using a voltage threshold of ±1000 *μV* and spectral threshold (range −100 dB to +30 dB) within the 20-40 Hz frequency band. Channels exhibiting artifacts in more than 20% of epochs were excluded from both the original and the copied dataset. ICA was then performed on the filtered copy and the resulting ICA weights were transferred back to the original dataset (Viola et al., 2010). Artifactual components were then automatically identified and removed using the Adjusted-ADJUST algorithm (Leach et al., 2020). The cleaned data were segmented into 1500 ms epochs, beginning 500 ms before stimulus onset. To further address residual artifacts, we applied a two-step rejection procedure (Morales et al., 2022). First, epochs were discarded if voltages from ocular electrodes (E1, E2, E5, E10, E11, and E17) exceeded ±125 *μV*. Second, for non-ocular channels, values exceeding ±150 *μV* were interpolated at the epoch level. If more than 10% of channels (excluding previously removed global channels) exceeded ±125 in an epoch, the entire epoch was rejected. Missing channels were interpolated using spherical spline interpolation and data were re-referenced to the average of the mastoid electrodes prior to quantification. For resting-state data, an average of 968.0 out of 1200 epochs (80.7%) were included for analyses. For oddball task data, an average of 767.5 out of 960 trials (80.0%) were included for analyses. Following quality control checks, 68 young and 66 older adults resting-state data and 54 young and 54 oddball task data were used for analyses.

### Aperiodic Activity

Full scalp EEG power spectra were parameterized using the Spectral Parametrization (specparam) toolbox in Python (Donoghue et al., 2020). As in prior work, we first visually inspected the nature of the aperiodic component of the power spectrum in log-log space and applied a 3-48 Hz frequency range with a resolution of 0.1 Hz resolution in fixed (i.e., linear) mode (Donoghue et al., 2020; Kopčanová et al., 2024). The following specparam toolbox settings were used: *peak width limits* (2.5-8), *peak threshold* (1.0), *aperiodic mode* (fixed), *maximum number of peaks* (6), and *minimum peak height* (0.05). Model fit quality was assessed using the mean *R^2^* value. The average goodness-of-fit of the final models averaged across tasks and electrodes was *R^2^* = 0.986 (SE = 0.048) for young adults, and *R^2^* = 0.980 (SE = 0.047) for older adults, demonstrating robust and comparable fits to those reported in previous studies with older adults (Donoghue et al., 2020; Kopčanová et al., 2024; Merkin et al., 2023). Given prior evidence of age-related changes in alpha power and shifts in individual peak alpha frequencies (Babiloni et al., 2006; Polich, 1997; Scally et al., 2018; Tröndle et al., 2023; Vysata et al., 2012), we first identified each participant’s individual peak alpha frequency (iPAF) by computing the center frequency within an extended alpha band range (5–15 Hz). Group-level mean iPAFs were then used to derive adjusted alpha frequency bands for young and older adults by shifting the band of older adults based on the mean group difference in iPAF. These group frequency ranges were then used to extract alpha band oscillatory power. In addition, the aperiodic exponent and offset were extracted for each participant and included in subsequent statistical analyses. Using the *specparam* toolbox (Donoghue et al., 2020), we confirmed that all young adults exhibited iPAFs in the typical alpha band frequency, 7-13 Hz, during eyes closed resting-state recordings. In contrast, older adults demonstrated a consistent slowing of iPAFs by 0.5 Hz across the frontal (Fz), central (Cz), and parietal (Pz) midline electrodes. Accordingly, we defined the alpha band as 7-13 Hz for young adults and 6.5-12.5 Hz for older adults when analyzing alpha power. To address our central research question concerning the impact of arousal on oscillatory alpha power, we used relative alpha power as expressed as a proportion of total spectral power across the other frequency bands (i.e., 3-48 Hz) to best conduct between-subjects comparisons.

### Event-Related Potential Analyses

P300 amplitude and peak latency were automatically extracted from stimulus-evoked ERP waveforms using custom MATLAB scripts using the EEGLAB toolbox (Delorme & Makeig, 2004). Baseline correction was applied using the −200 to 0 ms pre-stimulus window and ERPs were averaged across all artifact-free trials. In line with prior studies employing the auditory oddball paradigm (Polich, 2007; Walsh et al., 2017; Wronka et al., 2008), the P300 component was examined at Fz, Cz, and Pz midline electrodes in which P300 responses are most prominent. To define the temporal window of the P300 component, a data-driven approach was used. Specifically, we created a difference wave using the grand-averaged ERP waveforms for the passive and active oddball tasks, leveraging the fact that attentional engagement is required in the active but not the passive task. The time window was defined as the positive component above 0. The resulting difference waveforms were used to empirically define the P300 window separately for each group. For young adults, the P300 component was defined as the 246-390 ms post-stimulus window and for older adults, as the 288-432 post-stimulus window. P300 amplitudes were quantified by computing the area under the curve (AUC) within the respective time windows for each group. Peak latencies were determined using MATLAB’s *findpeaks* function to identify the maximum positive peak within the defined window, with the latency corresponding to the timing of that peak. All automatically identified peaks and latencies were visually inspected by two co-authors to ensure accuracy and consistency.

### Pupillometry Analyses

Eye-tracking data were collected during all resting-state and oddball task runs and stored in EDF format. Sample and event data were imported into MATLAB using the *edf2mex* MEX program. Pupil data were preprocessed using the automated artifact removal algorithm *ET-remove-artifacts* (Mather et al., 2020). In brief, this algorithm identifies blink-related artifacts by detecting abrupt changes in pupil velocity and linearly interpolates across the start and end points of each artifact to generate a cleaned time series. For each run, baseline pupil size was defined as the average pupil diameter across the entire recording period. Pupil dilation responses (PDRs) were quantified as the maximum change in pupil size relative to a 500 ms pre-stimulus baseline window. The PDR was defined as the maximal pupil dilation occurring within the 2000 ms window following stimulus onset. Following quality control checks, 63 young and 63 older adults’ pupillometry data were used for analyses. In addition, 67 young and 65 older adults’ behavioral data were used for analyses.

### Data Availability Statement

The behavior and eye tracking data have been made publicly available on the Open Science Framework, https://osf.io/xbcr5/. The raw EEG data have been made publicly available on OpenNeuro: Dataset 006466 for older adults, and 006480 for young adults.

## Results

### Arousal exerts age-dependent effects on response times to the distractor tone

We first investigated whether elevating arousal via threat of unpredictable shock differentially modulates behavioral response times across age groups. A 2 (Age: young, older) x 2 (Arousal: no-threat, threat) x 3 (Stimulus Type: standard, target, distractor) mixed ANOVA revealed a significant main effect of arousal, *F*(1,130) = 119.49, *p* < .001, *η_p_^2^* = .479, and a significant main effect of stimulus type, *F*(2,260) = 331.10, *p* < .001, *η_p_^2^* = .718. There was no main effect of age, *F*(1,130) = .00, *p* = .997. Critically, a significant three-way interaction emerged among age, arousal, and stimulus type, *F*(2,260) = 4.76, *p* = .009, *η_p_^2^* = .035. To further explore this interaction, we conducted separate 2 (Age) x 2 (Arousal) mixed ANOVAs for each stimulus type.

For standard tones, there was a significant main effect of arousal, *F*(1,130) = 60.34, *p* < .001, *η_p_^2^* = .317, with slower response times under arousal for both young (No-threat: *M =* 384.2 ms, *SE* = 99.0 ms; Threat: *M =* 415.1 ms, *SE* = 108.7 ms) and older adults (No-threat: *M =* 376.8 ms, *SE* = 74.0 ms; Threat: *M =* 410.0 ms, *SE* = 99.3 ms). No significant main effect of age, *F*(1,130) = .14, *p* = .700, nor age x arousal interaction, *F*(1,130) = .08, *p* = .782 was observed (see Figure 2). For target tones, there was again a significant main effect of arousal, *F*(1,130) = 104.34, *p* < .001, *η_p_^2^* = .445, with slower response times under arousal for both young (No-threat: *M =* 464.0 ms, *SE* = 108.4 ms; Threat: *M =* 522.0 ms, *SE* = 129.5 ms) and older adults (No-threat: *M =* 471.3 ms, *SE* = 90.2 ms; Threat: *M =* 512.7 ms, *SE* = 104.2 ms). No significant main effect of age, *F*(1,130) = .00, *p* = .954, nor age x arousal interaction, *F*(1,130) = 2.93, *p* = .089 was observed (see Figure 2). For distractor tones, a significant main effect of arousal was observed, *F*(1,130) = 57.74, *p* < .001, *η_p_^2^* = .308, along with a significant age x arousal interaction, *F*(1,130) = 8.24, *p* = .005, *η_p_^2^* = .060 (see Figure 2). Post-hoc *t*-tests revealed that elevated arousal significantly slowed response times for the distractor tone in young adults, *t*(66) = 6.45, *p* < .001, *d =* .393 (No-threat: *M* = 486.7 ms, *SE* = 101.6 ms; Threat: *M* = 539.2 ms, *SE* = 134.1 ms), but to a lesser extent in older adults as hypothesized, *t*(64) = 4.12, *p* < .001, *d* = .265 (No-threat: *M* = 508.3 ms, *SE* = 89.9 ms; Threat: *M* = 532.0 ms, *SE* = 89.2 ms). No main effect of age was found, *F*(1,130) = .16, *p* = .686. These findings indicate that the effect of arousal on older adults was specific to distractibility by salient but task-irrelevant stimuli.

**Figure 2.**
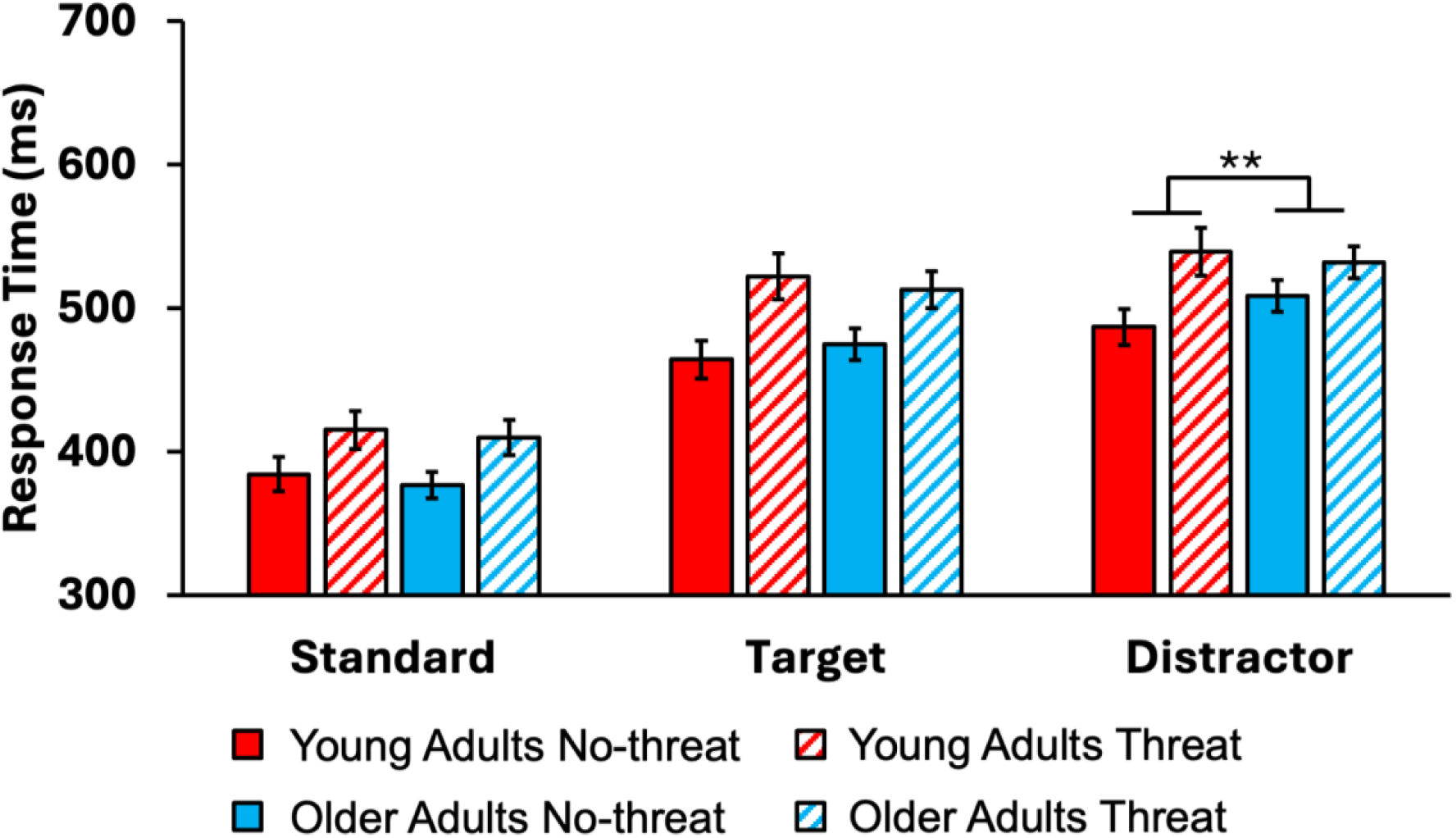
Elevated arousal significantly delayed response times in both young and older adults across all stimulus types, but the effect was attenuated only in older adults for distractor tones. Response times to standard tones reflect general responsiveness to frequent stimuli, target tone responses serve as an index of goal-directed attentional control, and distractor tone responses reflect involuntary attentional capture or distractibility by salient but task-irrelevant stimuli. A significant main effect of arousal was observed across all stimulus types, but a significant age x arousal interaction emerged specifically for distractor tones. Error bars reflect standard errors of the mean. \*\**p* < .01

### Older adults exhibit smaller evoked pupil responses in the active task

Pupillometry is the most widely used non-invasive method for indirectly assessing noradrenergic activity in humans. Pupil dilation responses have been reliably linked to phasic, task-evoked noradrenergic firing (Costa & Rudebeck, 2016; Elman et al., 2017; Gilzenrat et al., 2010; Joshi & Gold, 2020; Joshi et al., 2016; Liu et al., 2017; Murphy et al., 2011; Weigand et al., 2023), and inferences on tonic (baseline) noradrenergic firing are often drawn based on its inverse relationship with phasic responses (Aston-Jones & Cohen, 2005). In the rodent and non-human primate literature, LC neuronal firing has been directly linked to states of arousal and subsequent behavior (G. Aston-Jones & Bloom, 1981; G. Aston-Jones et al., 1999; Gary Aston-Jones & Cohen, 2005; Rajkowski et al., 1994a, 1994b; Sara & Bouret, 2012), characterized by tonic firing during relaxed states and phasic bursts during task engagement. By including both passive and active versions of the oddball task in this study, we examined these dynamics in humans and compared pupil responses in relaxed states in the passive task and during task-engaged, higher arousal states in the active task. Based on the hypothesis that aging is marked by elevated tonic noradrenergic activity, we predicted that older adults would exhibit reduced pupil dilation responses even in relaxed conditions. We further explored whether changes in arousal states would extend to phasic responses during active task engagement.

We first investigated whether aging differentially modulated evoked pupil responses under no-threat and threat conditions in the passive task. A 2 (Age: young, older) x 2 (Arousal: no-threat, threat) x 3 (Stimulus: standard, target, distractor) mixed ANOVA revealed a significant main effect of arousal, *F*(1,122) = 4.41, *p* = .038, *η_p_^2^*= .035, but no main effect of age, *F*(1,122) = 2.61, *p =* .109, nor stimulus*, F*(2,244) = 2.59, *p* = .084. In addition, we identified no significant interactions, *Fs*(2,244) < 1.49, *ps* > .229 (see Figure 3 and Table S1). In the active task, a mixed ANOVA revealed a significant main effect of age, *F*(1,124) = 6.10, *p* = .015, *η_p_^2^* = .047, and stimulus, *F*(2,248) = 38.62, *p* < .001, *η_p_^2^* = .237, but not arousal, *F*(1,124) = 2.54, *p* = .113. Importantly, there was a significant three-way age x arousal x stimulus interaction, *F*(2,248) = 3.23, *p* = .042, *η_p_^2^* = .025 (see Figure 3 and Table S1). To probe this interaction, we conducted separate age x arousal ANOVAs for each stimulus type. Significant main effects of age emerged for the standard and distractor tone, *Fs*(1,124) > 4.54, *ps* < .035, *η_p_^2^* > .035, but not for the target tone, *F*(1,124) 2.47, *p* = .119. In addition, we identified a significant main effect of arousal for the standard tone, *F*(1,124) = 10.05, *p* = .002, *η_p_^2^*= .075, but not for the target and distractor tones, *Fs*(1,124) < 1.77, *ps* > .185. No interactions were significant across any of the stimuli, *Fs*(1,124) < 2.00, *ps* > .160.

**Figure 3.**
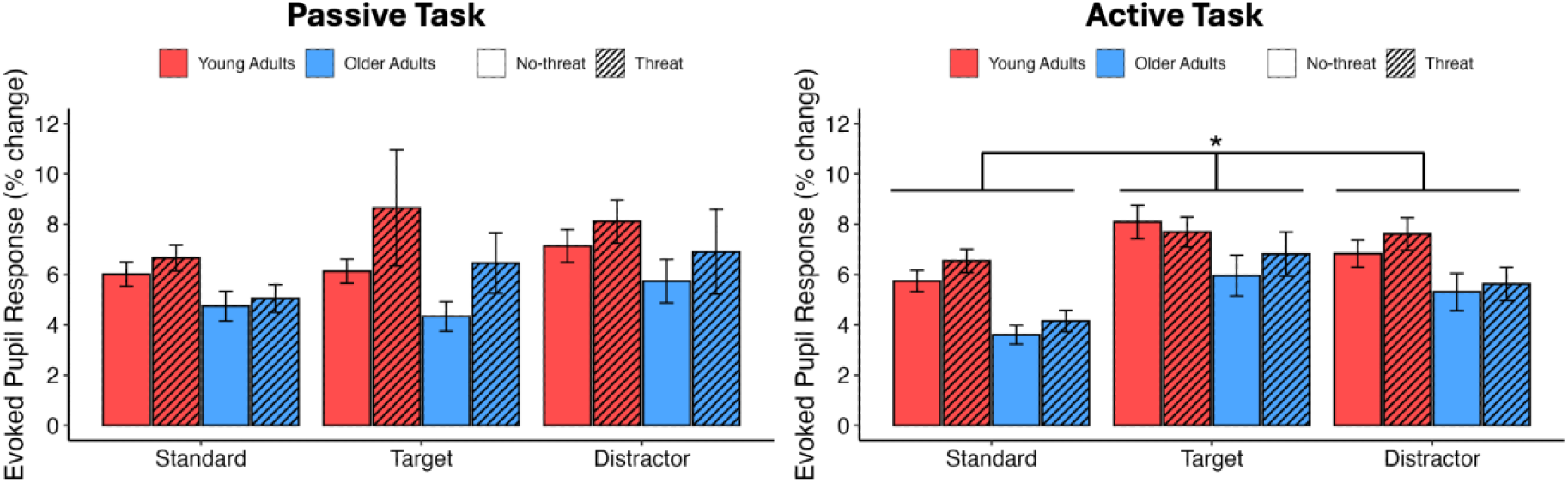
Pupillometry evidence for tonic hyperactivity in aging. Evoked pupil responses were calculated as percent change from baseline to account for age-related differences in absolute pupil size. (A) In the passive task, threat of shock increased pupil responses across both age groups. (B) In the active task, a significant three-way interaction of arousal, stimulus type, and age emerged. For both the standard and distractor tones, older adults exhibited smaller evoked pupil responses compared with young adults, but no age differences were found for the target tone. These findings indicate that during task engaged states, older adults show reduced phasic noradrenergic responses to repeated and task-irrelevant tones consistent with an elevated tonic noradrenergic state. However, responses to the target tone were age equivalent, suggesting that goal-directed processing evokes comparable responses across age groups. Error bars represent the standard error of the mean.

### Age differences in the P300 component during passive and active oddball tasks

Following seminal auditory oddball studies that have identified maximal P300 amplitudes along the midline scalp electrodes, (Mertens & Polich, 1997; Polich, 2007) we *a priori* selected frontal (Fz), central (Cz), and parietal (Pz) electrode sites for analysis. To evaluate whether age-related differences in P300 amplitude varied as a function of task, stimulus type, and scalp location, we conducted a 2 (Age: young, older) x 2 (Task: passive, active) x 3 (Stimulus: standard, target, distractor) x 3 (Electrode: Fz, Cz, Pz) mixed ANOVA. This analysis revealed a significant four-way interaction among age, task, stimulus type, and electrode, *F*(4,424) = 4.27, *p* = .002, *η_p_^2^*= .039, indicating that age-related differences in P300 amplitude were dependent on both task and scalp location (see Figure 4 for scalp topography maps). Given this interaction, we conducted follow-up analyses to examine age differences in P300 amplitudes separately at each electrode site.

**Figure 4.**
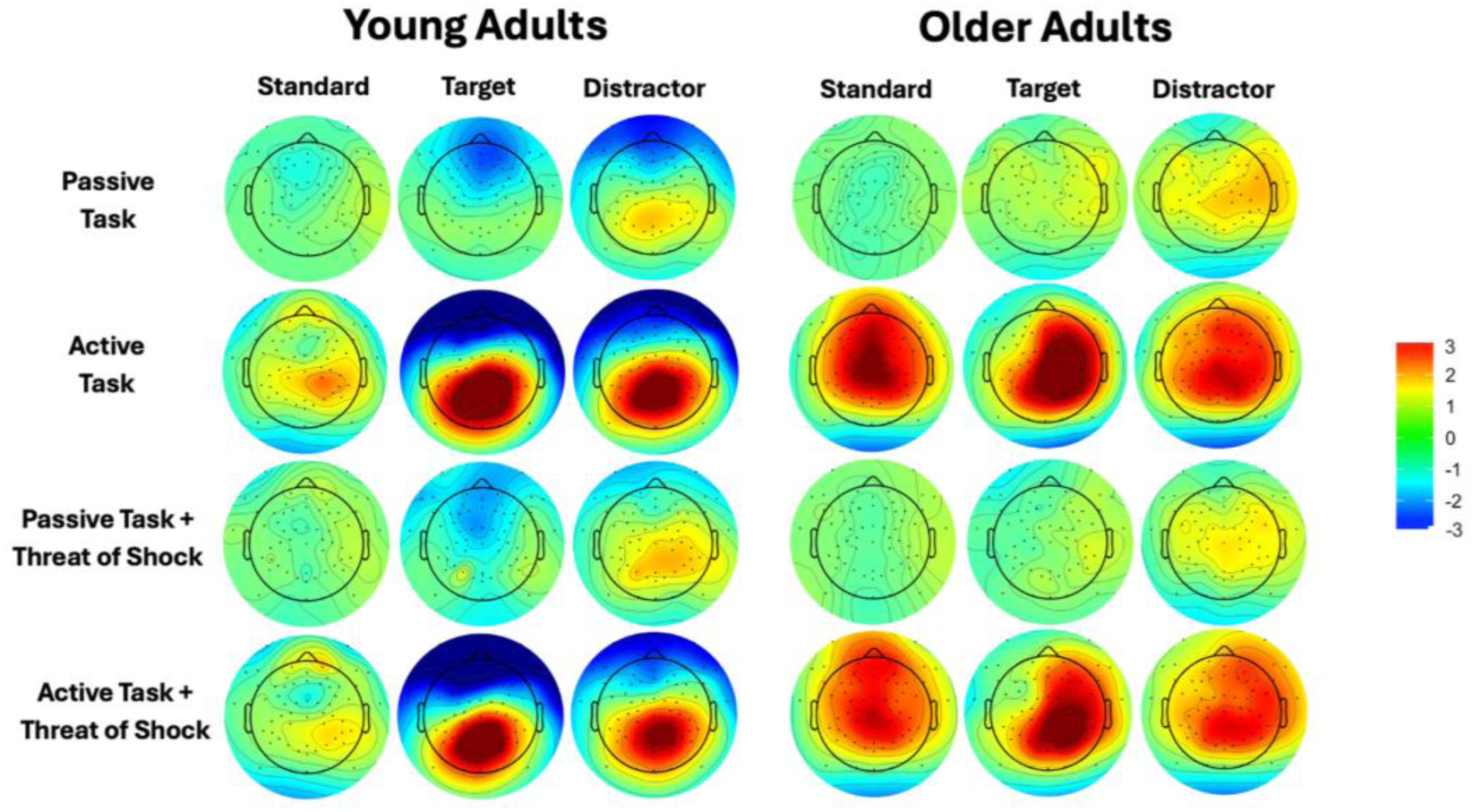
Scalp topography maps by age group, stimulus type, and task conditions. Topographic distributions of P300 amplitudes are displayed across young and older adults for each stimulus type (standard, target, distractor) and task conditions (passive, active, passive + threat of shock, active + threat of shock). Overlaid dots indicate electrode locations from the 65-channel Geodesic Sensor Net (GSN) used for EEG data acquisition.

### Frontal electrode findings suggest older adults exhibited greater attention processing

First, we examined age differences in frontal P300 responses at electrode Fz. With P300 amplitude as the dependent measure, a 2 (Age: young, older) x 2 (Task: passive, active) x 3 (Stimulus: standard, target, distractor) mixed ANOVA revealed significant main effects of age, *F*(1,106) = 20.73, *p* < .001, *η_p_^2^*= .164, task, *F*(1,106) = 22.09, *p* < .001, *η_p_^2^*= .172, and stimulus type, *F*(2,212) = 4.56, *p* = .012, *η_p_^2^*= .041. Importantly, a significant three-way interaction among age, task, and stimulus type was also observed, *F*(2,212) = 5.56, *p* = .004, *η_p_^2^*= .050 (see Figure 5A). Post hoc analyses indicated that in the active task, older adults exhibited significantly larger P300 amplitudes across all stimuli compared to the passive task, *ts*(53) *>* 3.28, *ps* < .002, *ds* .495. In contrast, young adults did not show significant differences in P300 amplitude between the passive and active conditions, *ts* < 1.57, *ps* > .123 (see Table S2). Moreover, in the active task, older adults demonstrated significantly greater P300 amplitudes than young adults for all stimulus types, *ts*(106) > 3.06, *ps* < .003, *ds* > .590. This pattern was also evident in the passive task for the target and distractor tones, *ts*(106) > 2.95, *ps* < .004, *ds* > .568, although no age difference was observed for standard tones, *t*(106) = .23, *p* = .817. These findings suggest that older adults exhibit greater allocation of frontal resources compared with young adults, but alternatively may reflect age differences in the underlying P300 dipole scalp localization.

**Figure 5.**
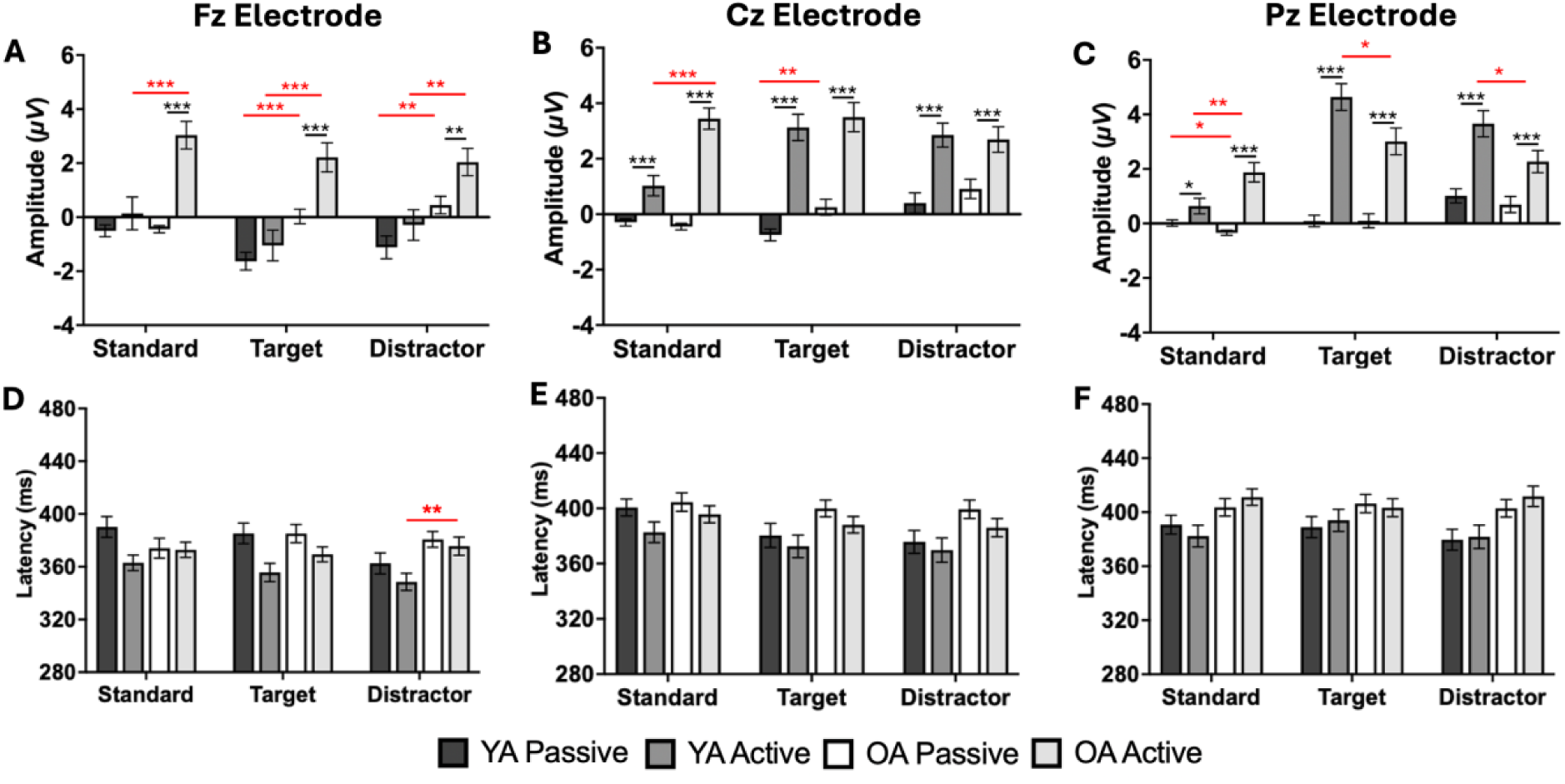
Age differences in the P300 component while completing the passive and active auditory oddball tasks. (A) When task engaged, older adults showed stronger P300 responses in the frontal electrode compared with young adults. (B) In the central electrode, young adults showed the typical pattern of greater P300 amplitudes to the target and distractor tone compared with the standard tone. However, older adults showed no significant differences in P300 amplitudes across tone types. (C) Finally, older adults showed significantly smaller P300 amplitudes compared with young adults to the target and distractor tone over the parietal Pz electrode, unlike in the central electrode with age-equivalent amplitudes. However, older adults still showed greater P300 amplitudes than young adults for the standard tone in the active task, as also seen in the central electrode. (D-F) Over all electrode sites, older adults also showed slower P300 peak latencies compared with young adults. YA = young adults. OA = older adults.

When examining P300 peak latency at Fz, we did not observe a significant main effect of age, *F*(1,106) = 1.89, *p* = .172. However, there were significant main effects of task, *F*(1,106) = 7.80, *p* = .006, *η_p_^2^*= .069, and stimulus type, *F*(2,212) = 4.44, *p* = .013, *η_p_^2^*= .040. In addition, we observed a significant stimuli x age interaction, *F*(1,106) = 9.59, *p* < .001, *η_p_^2^*= .083 (see Figure 5D). Post hoc comparisons revealed that, in the active task, older adults exhibited significantly slower P300 peak latencies than young adults in response to distractor tones, *t*(106) = 2.87, *p* = .005, *d* = .553, but no significant age differences were found for standard or target tones, *ts*(106) < 1.83, *ps >* .070.

### Older adults exhibit heightened central processing of the standard tone leading to reduced stimulus differentiation

At the central electrode (Cz), a 2 (Age: young, older) x 2 (Task: passive, active) x 3 (Stimulus: standard, target, distractor) mixed ANOVA on P300 amplitudes revealed no significant main effect of age, *F*(1,106) = 3.34, *p* = .071. However, we observed significant main effects of task, *F*(1,106) = 106.89, *p* < .001, *η_p_^2^*= .502, stimulus type, *F*(2,212) = 11.77, *p* < .001, *η_p_^2^*= .100, and a significant three-way interaction among age, task, and stimulus, *F*(2,212) = 24.51, *p* < .001, *η_p_^2^*= .188 (see Figure 5B). Both young and older adults demonstrated significantly greater P300 amplitudes in the active task compared with the passive task across all stimuli, *ts*(53) > 4.08, *ps* < .001, *ds* > .567. In the passive task, older adults showed significantly larger P300 amplitudes than young adults in response to the target tone, *t*(106) = 2.80, *p =* .006, *d* = .538, but no group differences were observed for the standard or distractor tones, *ts* < 1.02, *ps* > .156. In the active task, older adults exhibited significantly larger P300 amplitudes than young adults for the standard tone, *t*(106) = 4.57, *p <* .001, *d* = .880, but not for the target or distractor tones, *ts* < .53, *ps* > .599. As typically observed in the auditory oddball paradigm, young adults showed significantly larger P300 amplitudes to the target tone relative to the standard tone, *t*(53) = 6.45, *p* < .001, *d = .*646. However, older adults exhibited no such differentiation, showing comparable P300 amplitudes to the standard and target tones, *t*(53) = .17, *p* = .869, again suggesting that older adults allocated increased attentional resources to the frequent stimulus leading to diminished differentiation between the standard and target tones. Interestingly, in the passive task, young adults showed reduced P300 amplitudes to the target compared with the standard tone, *t*(53) = 2.33, *p* = .024, *d* = .330, while older adults showed the opposite pattern with larger P300 amplitudes to the target tone, *t*(53) = 2.70, *p* = .009, *d* = .399.

When examining P300 peak latency at Cz, a significant main effect of age, *F*(1,106) = 4.21, *p* = .043, *η_p_^2^*= .038, and stimuli was observed, *F*(2,212) = 13.46, *p* < .001, *η_p_^2^*= .113, but no significant main effect of task, *F*(1,106) = 3.82, *p* = .053 (see Figure 5E).

### Older adults exhibit Posterior-to-Anterior Shift (PASA) in aging

At the parietal electrode (Pz), a 2 (Age: young, older) x 2 (Task: passive, active) x 3 (Stimulus: standard, target, distractor) mixed ANOVA on P300 amplitude revealed no significant main effect of age, *F*(1,106) = 1.51, *p* = .222. However, significant main effects were observed for task, *F*(1,106) = 94.17, *p* < .001, *η_p_^2^*= .470, stimulus type, *F*(2,212) = 53.77, *p* < .001, *η_p_^2^*= .337, and three-way interaction among age, task, and stimulus, *F*(2,212) = 20.55, *p* < .001, *η_p_^2^*= .162 (see Figure 5B). Consistent with results at Cz, both young and older adults showed significantly greater P300 amplitudes during the active task compared with the passive task across all stimuli as expected, *ts*(53) > 2.10, *ps* < .041, *ds* > .381. While older adults showed greater P300 amplitudes at Cz compared with young adults, the opposite pattern emerged at Pz during the active task. Young adults exhibited larger P300 amplitudes than older adults for both the target and distractor tones, *ts*(106) > 2.21, *ps* < .029, *ds* > .425. Interestingly, for the standard tone, older adults continued to show greater amplitudes than young adults, *t*(106) = 2.71, *p* = .008, *d* = .521. For young adults, P300 amplitude was largest for the target tone compared to both the standard, *t*(53) = 10.70, *p* < .001, *d* = 1.22, and distractor tones, *t*(53) = 3.00, *p* = .004, *d* = .275. Similarly, distractor tones elicited larger P300 amplitudes than standard tones, *t*(53) = 8.63, *p* < .001, *d* = .921. Among older adults, the target tone also evoked the largest P300 response relative to standard and distractor tones, *ts*(53) > 2.77, *ps* < .008, *ds* .214. However, no significant difference was observed between the target and distractor tones, *t*(53) = 1.51, *p* = .136.

When examining P300 peak latency at Pz, we identified a significant main effect of age, *F*(1,106) = 7.56, *p* = .007, *η_p_^2^*= .067, but no significant main effects of task, *F*(1,106) = .12, *p* = .728, stimuli, *F*(2,212) = 1.10, *p* = .336, nor three-way interaction, *F*(2,212) = 2.91, *p* = .057 (see Figure 5E).

### Age differences in the effect of arousal on the P300 Component

To investigate whether arousal differentially modulated P300 responses across age groups, we conducted a 2 (Age: young, older) x 2 (Task: passive, active) x 2 (Arousal: no-threat, threat) x 3 (Stimulus: standard, target, distractor) mixed ANOVA. In this section, we report only the effects related to arousal and its interactions, to avoid redundancy.

At the Fz electrode, there was no significant main effect of arousal on P300 amplitude, *F*(1,106) = 1.97, *p* = .163. However, a significant task x arousal interaction emerged, *F*(1,106) = 12.61, *p* < .001, *η_p_^2^*= .106. Post hoc comparisons revealed that in the passive task, arousal significantly increased P300 amplitudes to the distractor tone in young adults, *t*(53) = 2.32, *p* = .024, *d* = .343. In the active task, arousal reduced P300 amplitudes to the target tone in young adults, *t*(53) = 3.25, *p =* .002, *d* = .268, and to the standard tone in older adults, *t*(53) = 2.50*, p* = .016, *d* = .164. With P300 peak latencies as the dependent measure, no significant main effect of arousal was found, *F*(1,106) = 1.72, *p* = .192, and no significant interactions were observed.

**Figure 6.**
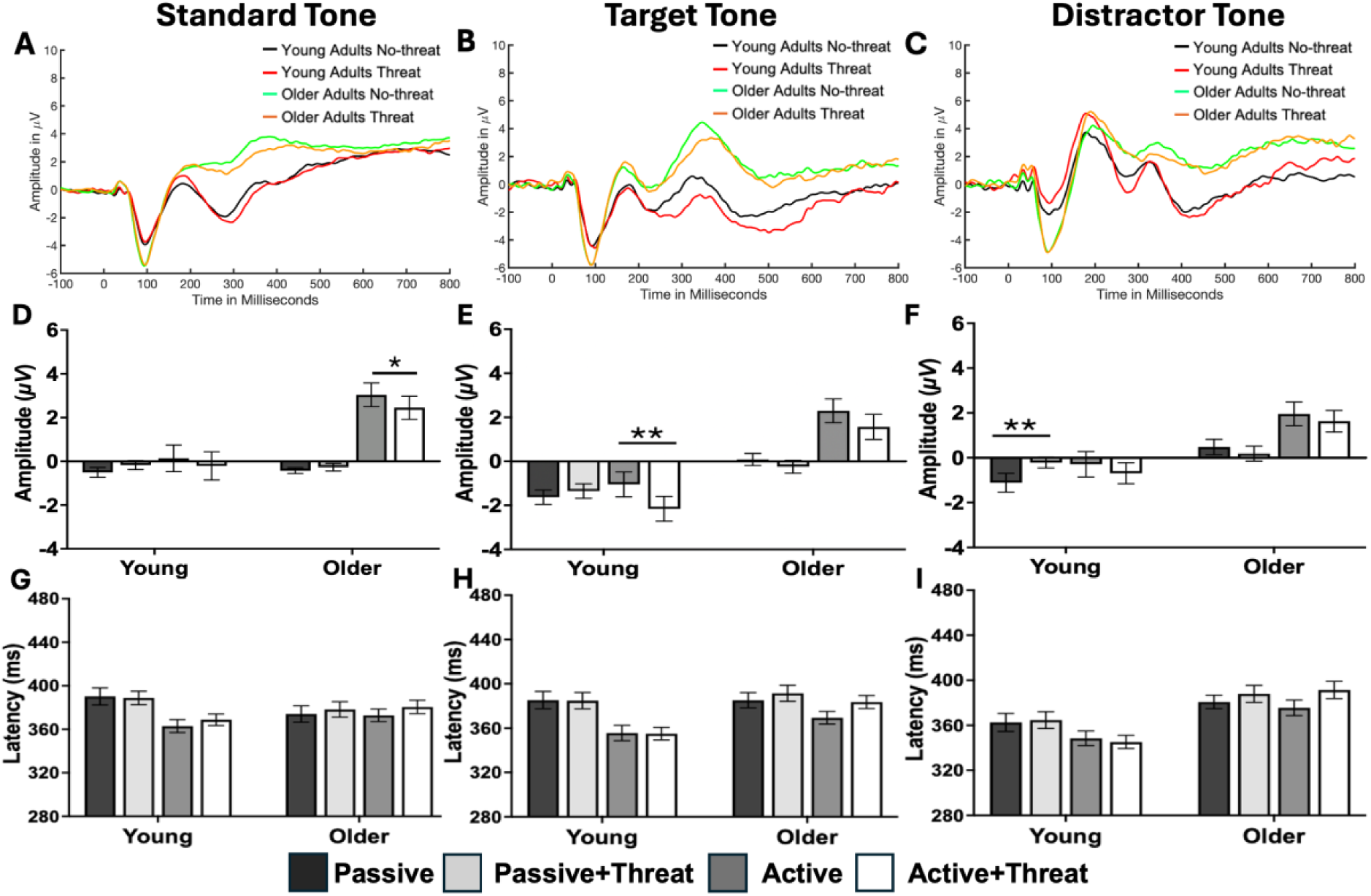
In the Fz electrode, arousal increased amplitudes in the passive task for young adults but decreased amplitudes in the active task. Panels A-C display the grand average ERP waveforms for the (A) standard, (B) target, and (C) distractor tones across both age groups under no-threat and threat conditions. Panels D-F show mean P300 amplitudes for the (D) standard, (E) target, and (F) distractor tones, and reveal a significant task x arousal interaction with arousal increasing P300 amplitudes in the passive task but decreasing them in the active task. Panels G-I present mean P300 peak latencies for the (G) standard, (H) target, and (I) distractor tones. No significant effects of arousal were observed for peak latencies. Error bars represent the standard error of the mean.

At the Cz electrode, analysis of P300 amplitudes revealed a significant main effect of arousal, *F*(1,106) = 13.06, *p* < .001, *η_p_^2^*= .110, in addition to a significant task x arousal interaction, *F*(1,106) = 19.96, *p* < .001, *η_p_^2^*= .158, and a significant arousal x stimulus interaction, *F*(2,212) = 4.58, *p* = .011, *η_p_^2^*= .041. In the active task, arousal significantly reduced P300 amplitudes across all stimulus types for both young and older adults, *ts*(53) > 2.30, *ps* < .025, *ds* > .219, except for the distractor tone in older adults, *t*(53) = 1.78, *p* = .081. However, in the passive task, arousal had no significant effect on P300 amplitudes, *ts*(53) < 1.90, *ps* .063. With P300 peak latencies as the dependent measure, we observed a significant main effect of arousal, *F*(1,106) = 8.31, *p* = .005, *η_p_^2^*= .073, in addition to a significant three-way interaction between age x task x arousal, *F*(1,106) = 5.36, *p* = .023, *η_p_^2^*= .048, and between age x arousal x stimulus type, *F*(1,106) = 3.23, *p* = .041, *η_p_^2^*= .030. In the active task, arousal delayed P300 peak latencies across all stimuli in older adults, *ts*(53) > 2.58, *ps* < .013, *ds* .415. In contrast, arousal did not significantly affect peak latencies in young adults, *ts(53)* < 1.00, *ps* > .322. In the passive task, arousal did not significantly modulate peak latencies for older adults, *ts*(53) < 1.07, *ps* > .291. However, among young adults, arousal significantly delayed peak latencies for the target tone, t(53) = 1.78, *p* = .012, *ds* = .406, but not for the standard tone, *t*(53) = 1.78, *p* = .082, or the distractor tone, *t*(53) = 1.36, *p* = .180.

**Figure 7.**
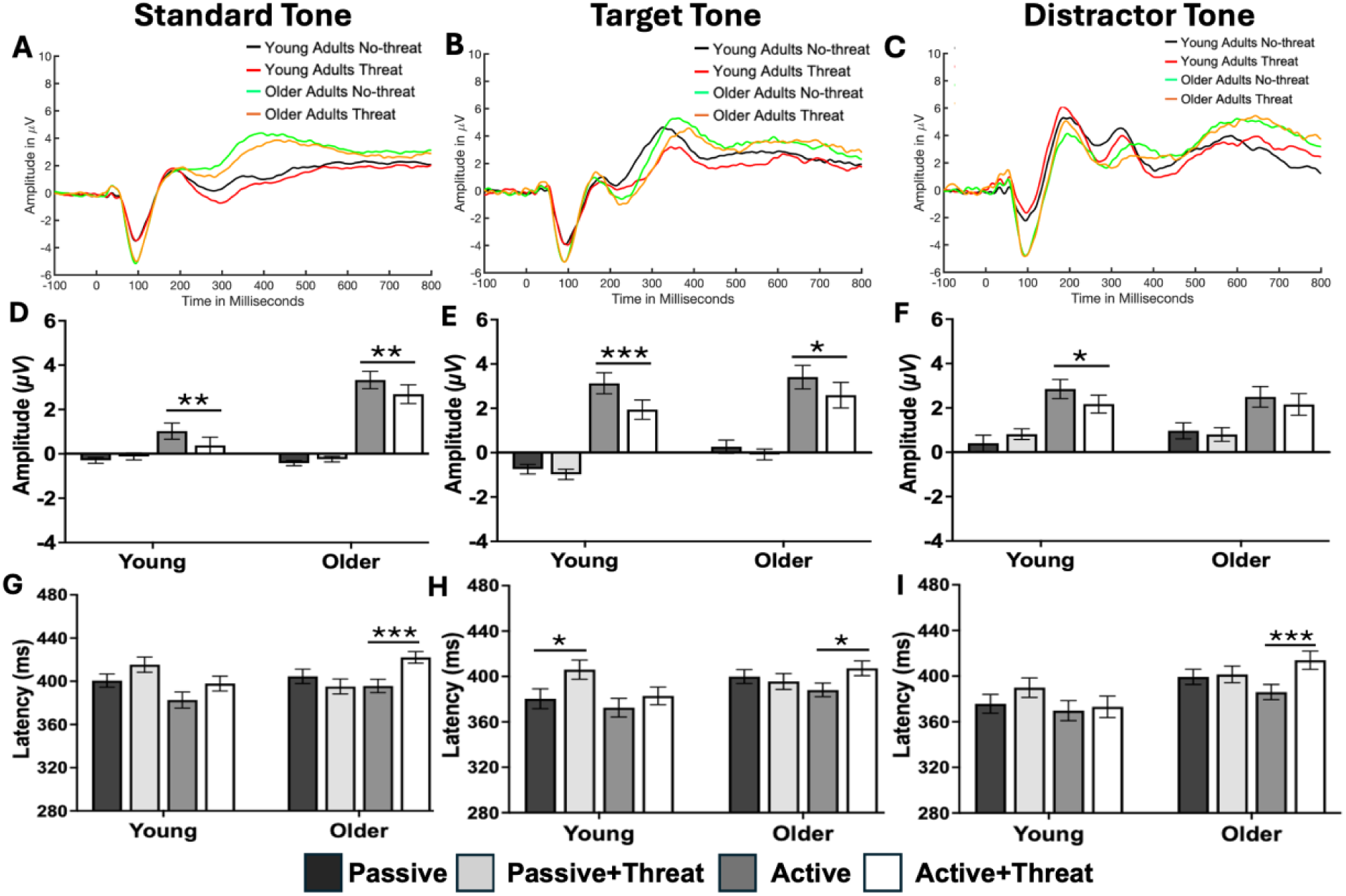
In the Cz electrode, arousal decreases amplitudes in the active task and delays latencies in both tasks. Panels A-C display the grand average ERP waveforms for the (A) standard, (B) target, and (C) distractor tones across both age groups under no-threat and threat conditions. Panels D-F show mean P300 amplitudes for the (D) standard, (E) target, and (F) distractor tones, and reveal that arousal significantly reduced amplitudes in the active task for both young and older adults. This arousal effect was not observed in the passive task. Panels G-I present mean P300 peak latencies for the (G) standard, (H) target, and (I) distractor tones, highlighting a significant age x arousal x stimulus interaction. Arousal delayed peak latencies in older adults during the active task across stimuli, but not in young adults. Conversely, in the passive task, arousal delayed peak latency for the target tone in young adults. Error bars represent the standard error of the mean.

At the Pz electrode, analysis of P300 amplitudes revealed no significant main effect of arousal, *F*(1,106) = 3.16, *p* = .078, but showed a significant age x arousal interaction, *F*(1,106) = 5.11, *p* = .026, *η_p_^2^*= .046. In the passive task, arousal significantly reduced P300 amplitudes in young adults for both the standard and target tones, *ts*(53) > 2.21, *ps* < .031, *ds* < .400. However, arousal had no effect on P300 amplitudes in older adults across any stimulus type, *ts*(53) < .61, *ps* > .547. For P300 peak latencies, a significant main effect of arousal was observed, *F*(1,106) = 4.44, *p* = .037, *η_p_^2^*= .040, with no significant interaction effects.

**Figure 8.**
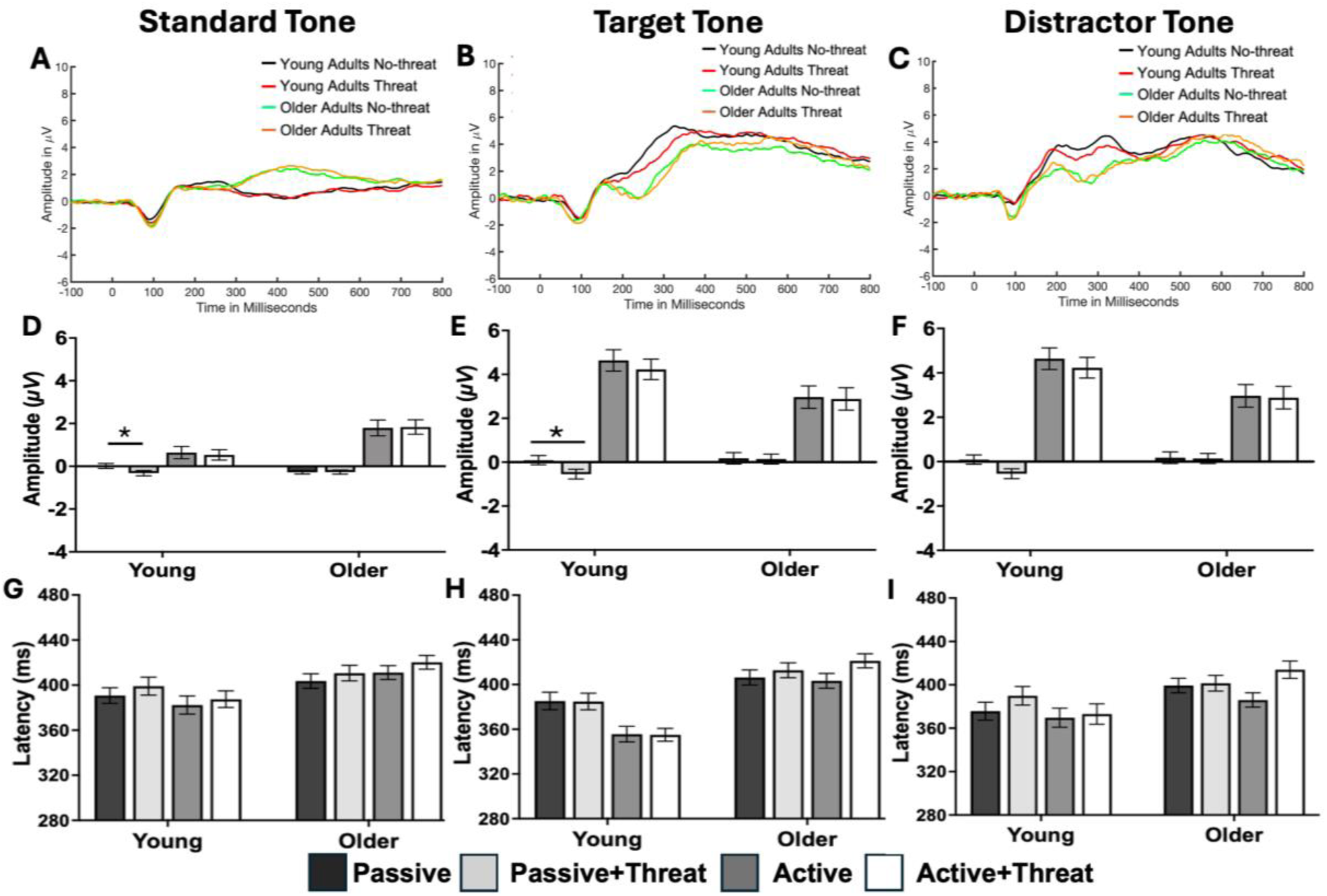
In the Pz electrode, arousal decreases amplitudes in the passive task for young adults. Panels A-C display the grand average ERP waveforms for the (A) standard, (B) target, and (C) distractor tones across both age groups under no-threat and threat conditions. Panels D-F show mean P300 amplitudes for the (D) standard, (E) target, and (F) distractor tones, and reveal that arousal significantly reduced amplitudes in young adults during the passive task for the standard and target tones, but had no effect in older adults. Panels G-I present mean P300 peak latencies for the (G) standard, (H) target, and (I) distractor tones, showing a significant main effect of arousal across conditions, with no significant age or task interactions. Error bars represent the standard error of the mean.

### Arousal differentially effects alpha power in the frontal and central electrodes across age groups

We examined whether increased arousal modulated resting-state EEG differently by age through a 2 (Age: young, older) x 2 (Condition: eyes closed, eyes open) x 2 (Arousal: no-threat, threat) x 3 (Electrode: Fz, Cz, Pz) mixed ANOVA with relative alpha power as the dependent measure and identified a significant four-way interaction, *F*(2,264) = 4.64, *p* = .010, *η_p_^2^*= .034. Thus, we again elected to explore the effect of arousal at each electrode location.

At the Fz electrode, we observed significant main effects of age, *F*(1,132) = 22.10, *p* < .001, *η_p_^2^*= .143, condition, *F*(1,132) = 156.39, *p* < .001, *η_p_^2^*= .542, arousal, *F*(1,132) = 12.48, *p* < .001, *η_p_^2^*= .086, and a significant three-way interaction of age x condition x arousal, *F*(1,132) = 8.77, *p* = .004, *η_p_^2^*= .062. At the Cz electrode, we again observed significant main effects of age, *F*(1,132) = 20.34, *p* < .001, *η_p_^2^*= .134, condition, *F*(1,132) = 141.97, *p* < .001, *η_p_^2^*= .518, arousal, *F*(1,132) = 32.99, *p* < .001, *η_p_^2^*= .200, and a significant three-way interaction, *F*(1,132) = 5.86, *p* = .017, *η_p_^2^*= .042. At the Pz electrode, we observed significant main effects of age, *F*(1,132) = 22.79, *p* < .001, *η_p_^2^*= .147, condition, *F*(1,132) = 95.19, *p* < .001, *η_p_^2^*= .419, and arousal, *F*(1,132) = 45.66, *p* < .001, *η_p_^2^*= .257. Additionally, there was a significant condition x arousal interaction, *F*(1,132) = 58.31, *p* < .001, *η_p_^2^*= .306, but no three-way interaction, *F*(1,132) = 1.84, *p* = .177. As expected, both young and older adults exhibited greater alpha power during eyes-closed compared with the eyes-open condition, *ts*(65) > 7.26, *ps* < .001, *ds* > .686. In addition, young adults demonstrated greater alpha power than older adults across both conditions and all electrode sites, *ts*(132) > 3.57, *ps* < .001, *ds* > .616.

To further isolate the effects of arousal, we post-hoc conducted separate 2 (Age: young, older) x 2 (Condition: no-threat, threat) mixed ANOVAs for the eyes-closed and eyes-open conditions. In the eyes-open condition, we observed robust main effects of arousal at all three electrode sites, *Fs*(1,132) > 23.29, *ps* < .001, *η_p_^2^s* > .150, but no significant age x arousal interactions, *Fs*(1,132) < 1.37, *ps* > .244. Arousal significantly increased relative alpha power in both young and older adults across all electrodes, *ts*(65) > 2.14, *ps* < .036, *ds* > .155. In the eyes closed condition, we did not observe main effects of arousal, *Fs*(1,132) < 3.44, *ps* > .066, but identified significant age x arousal interactions at the Fz and Cz electrodes, *Fs*(1,132) > 4,84, *ps* < .030, *η_p_^2^* > .035, but not at Pz, *F*(1,132) = .43, *p* = .514. Specifically, arousal led to increased alpha power at Fz and Cz in older adults, *ts*(65) > 2.14, *ps* < .036, *ds* > .101, but not at Pz, *t*(65) = 1.96, *p* = .054. No significant arousal-related changes in alpha power were observed in young adults during the eyes-closed conditions, *ts*(67) < 1.52, *ps* > .134.

**Figure 9.**
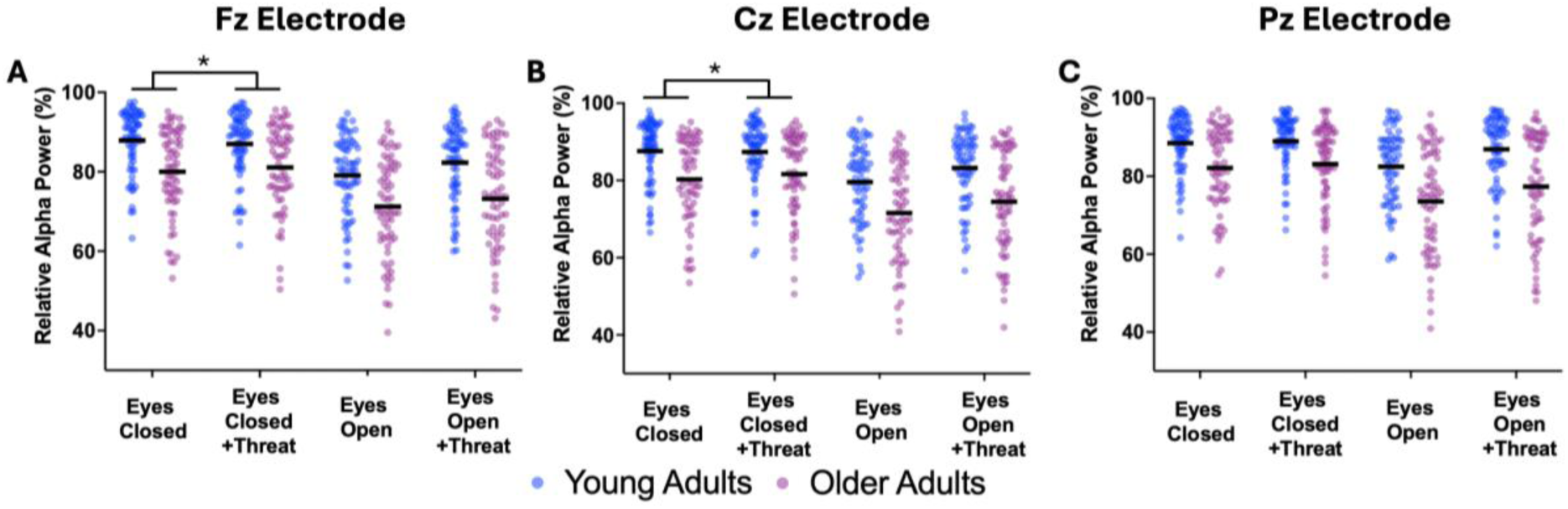
Arousal increases relative alpha power for older adults but not young adults in the Fz and Cz electrodes during eyes closed conditions. In the eyes closed condition, we identified a significant age x arousal interaction over the (A) Fz and (B) Cz electrodes, but not over the (C) Pz electrode. \**p < .*05

### Arousal exhibited age-equivalent effects on measures of aperiodic activity

We next examined age-related differences in aperiodic exponent and offset values across the Fz, Cz, and Pz electrodes in separate 2 (age: young, older) x 2 (condition, eyes closed, eyes open) x 2 (condition: no-threat, threat) x 3 (electrode: Fz, Cz, Pz) mixed ANOVA analyses. For the aperiodic exponent, we identified significant main effects of age, *F*(1,132) = 22.52, *p* < .001, *η_p_^2^*= .146, condition, *F*(1,132) = 60.18, *p* < .001, *η_p_^2^*= .313, arousal, *F*(1,132) = 4.86, *p* = .029, *η_p_^2^*= .035, and electrode, *F*(2,264) = 8.89, *p* < .001, *η_p_^2^*= .063. We also identified significant age x electrode, *F*(2,264) = 5.79, *p* = .003, *η_p_^2^*= .042, and arousal x electrode interactions, *F*(2,264) = 3.97, *p* = .020, *η_p_^2^*= .029. At the Cz electrode, arousal increased aperiodic exponent in eyes-closed condition for young adults, *t*(67) = 2.12, *p* = .038, *d* = 180. At the Pz electrode, arousal increased aperiodic exponent in the eyes-closed condition for older adults, *t*(65) = 2.11, *p* = .039, *d* = .201. For the aperiodic offset, we identified significant main effects of age, *F*(1,132) = 5.09, *p* = .026, *η_p_^2^*= .037, condition, *F*(1,132) = 63.45, *p* < .001, *η_p_^2^*= .325, arousal, *F*(1,132) = 29.25, *p* < .001, *η_p_^2^*= .181, and electrode, *F*(2,264) = 100.79, *p* < .001, *η_p_^2^*= .433. In addition, we identified a significant arousal x electrode interaction, *F*(2,264) = 4.69, *p* = .010, *η_p_^2^*= .034. At the Fz electrode, arousal did not modulate aperiodic offset in young adults across both conditions, *ts*(67) < 1.63, *ps* > .109, or in older adults during the eyes-open condition, *t*(65) = 1.55, *p* = .127. However, arousal increased aperiodic offset in the eyes-closed condition for older adults, *t*(65) = 3.64, *p* < .001, *d* = .231. At the Cz electrode, arousal increased aperiodic offset in both conditions for young adults, *t*(67) > 2.09, *ps* < .040, *ds* > .178, and older adults, *ts*(65) > 3.24, *ps* < .002, *ds* > .226. At the Pz electrode, arousal increased aperiodic offset in the eyes-open condition for young adults, *t*(67) = 2.77, *p* = .007, *d* = .236, and in both conditions for older adults, *ts*(65) > 3.19, *ps* < .002, *ds* > .273. Arousal did not affect aperiodic offset in the eyes-closed condition for young adults, *ts*(67) = .95, *p* = .344.

**Figure 10.**
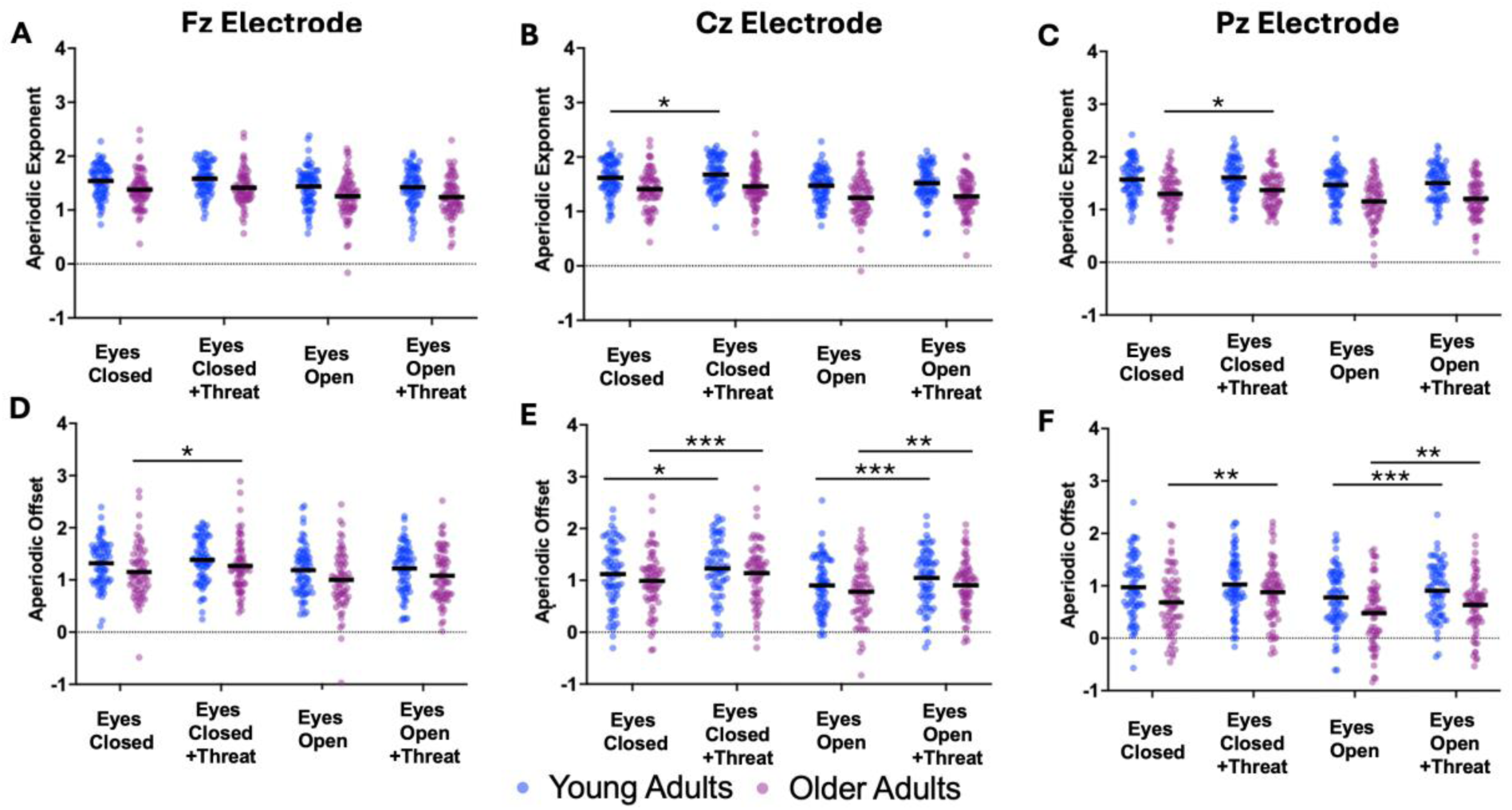
Arousal did not differentially modulate measures of aperiodic activity across age groups. We extracted aperiodic (A-C) exponent and (D-F) offset values at the Fz, Cz, and Pz electrodes. In the eyes-closed condition, arousal significantly increased the aperiodic exponent in young adults at the Cz electrode (B) and in older adults at the Pz electrode (C). For aperiodic offset, we identified a significant arousal x electrode interaction.

### Arousal differentially modulates behavioral response times and P300 latencies to the salient distractor in young and older adults

Finally, we examined whether the effect of arousal differed between young and older adults across six dependent measures: behavioral response time (RT), pupil dilation responses, alpha power, P300 amplitude, P300 peak latency, and aperiodic exponent. For the four task-related measures (RT, PDR, P300 amplitude and latency), we explored age differences in response to the distractor tone given as significant age x arousal interactions emerged for only this stimulus. For the spectral measures (alpha power, aperiodic exponent), we extracted periodic and aperiodic activity from the Fz electrode given that the P3a elicited by the distractor tone has a frontal scalp distribution (Mertens & Polich, 1997; Polich, 2007; Wronka et al., 2008, 2012). We fit a linear mixed-effects model with fixed effects of age group (young vs. older), modality (i.e., RT, PDR, alpha power, P300 amplitude, P300 latency, aperiodic exponent), and their interaction, as well as a random intercept for each participant in R (version 4.5). A Type III Satterwaite-corrected ANOVA (given unbalanced samples across modalities) revealed no main effect of age *F*(1, 736) = .26, *p* = .61, indicating that the overall magnitude of the arousal effect did not differ between age groups when averaged across modalities. However, there was a robust main effect of modality, *F*(5, 736) = 27.09, *p* < .001, *η_p_^2^*= .155, indicating that the impact of arousal varied significantly across the six measures. Critically, we observed a significant age x modality interaction, *F*(5, 736) = 6.54, *p* < .001, *η_p_^2^*= .043, indicating that age-related differences were modality specific. To further probe this interaction, we conducted Bonferroni-corrected pairwise comparisons of estimated marginal mean within each modality. Significant age differences emerged for behavioral response times, *t*(684) = 4.90, *p* < .001, *d* = .81, with young adults exhibiting increased slowing under threat of shock (*M* = 52.48 ms, *SE* = 4.40, 95% CI [44.38 60.57]) than older adults (*M* = 23.68 ms, *SE* = 4.50, 95% CI [15.48, 31.92]), and P300 latency, *t*(682) = -2.95, *p* = .003, *d* = .54, with older adults showing greater P300 delays (*M* = 15.83 ms, *SE* = 4.59, 95% CI[6.81, 24.85]) than young adults (*M* = -3.33 ms, *SE* = 4.59, 95% CI[-12.35, 5.69]). No significant age differences were observed for the remaining modalities, *ts* < .34, *ps* > .736. These findings show that increased arousal differentially modulates attentional control to salient but task-irrelevant distractors between young and older adults, specifically influencing attention processing speeds as observed in behavioral response times and event-related potentials (P300 component).

## Discussion

We identified converging evidence for LC-NA system hyperactivity in aging across multiple psychophysiological modalities including behavior, event-related potentials, and resting-state oscillations (see Table 1 for a summary). First, increased arousal led to significantly smaller behavioral slowing in response to the distractor tone among older adults compared with young adults, supporting the hypothesis of diminished arousal effects in older adults due to elevated tonic (i.e., baseline) LC activity. Second, pupil dilation responses during the active task were attenuated in older adults for the standard and distractor tone, consistent with a heightened and suboptimal tonic noradrenergic state that suppresses phasic responses (Aston-Jones & Cohen, 2005; Hayat et al., 2020). Finally, linear mixed model analyses showed significantly delayed P300 latencies in older adults compared with young adults under threat of shock. Although pharmacological and lesion studies have highlighted LC-NA contributions to P300 amplitude, some studies have reported mixed findings that show LC pharmacological antagonists lead to both faster and slower P300 latencies (Nieuwenhuis et al., 2005), which primarily index stimulus evaluation time (Magliero et al., 1984; Polich, 1987, 2007; Walsh et al., 2017). Our findings suggest that LC tonic hyperactivity alters neuromodulatory mechanisms of stimulus evaluation that leads to delayed P300 latencies. Prior studies have also shown that high-arousal words disrupts stimulus processing in an emotional Stroop task and leads to slower response times (Imbir et al., 2017), while low-arousal stimuli enhance evaluation speed (Purkis et al., 2009). Furthermore, rodent studies also show that auditory stimulus encoding is optimal at moderate arousal, whereas both high and low arousal conditions impairs stimulus discriminability (Papadopoulos et al., 2024). These findings support our interpretation that delayed P300 latencies observed in our study results is a consequence of suboptimal arousal states, and that this interaction is exacerbated in older adults who sustain tonic LC hyperactivity compared with young adults. Consistent with our findings, previous research has demonstrated that LC phasic response latency closely tracks manual response times on a trial-to-trial basis (Bouret & Sara, 2004), underscoring the coupling between these physiological modalities. Collectively, our behavioral, pupillometric, and electrophysiological data provide converging evidence that cognitively healthy older adults sustain elevated tonic noradrenergic activity compared with young adults.

**Table 1.** Summary of aging, arousal, and interaction effects by modality. Effects reflect results from the active task and the eyes-closed condition for resting state measures. Main effects of aging and arousal are interpreted relative to older adults (e.g., “decrease” indicates a reduced effect in older adults). In addition, main effects of arousal are interpreted relative to no-threat conditions (e.g., “decrease” indicates a reduced effect in threat conditions relative to no-threat conditions). For interaction effects, “OA < YA” indicates that the arousal effect was smaller in older adults compared with young adults. “--” indicates no significant statistical change.

| Measure | Electrode | Stimulus | Aging | Arousal | Interaction |
| --- | --- | --- | --- | --- | --- |
| Response Time |  | Standard | -- | increase | -- |
|  |  | Target | -- | increase | -- |
|  |  | Distractor | -- | increase | OA < YA |
| Pupil Dilation Response |  | Standard | decrease | increase | -- |
|  |  | Target | -- | -- | -- |
|  |  | Distractor | decrease | -- | -- |
| P300 Amplitude | Fz | Standard | increase | decrease | -- |
|  |  | Target | increase | -- | -- |
|  |  | Distractor | increase | -- | -- |
|  | Cz | Standard | increase | decrease | -- |
|  |  | Target | -- | decrease | -- |
|  |  | Distractor | -- | -- | -- |
|  | Pz | Standard | increase | -- | -- |
|  |  | Target | decrease | -- | -- |
|  |  | Distractor | decrease | -- | -- |
| P300 Latency | Fz | Standard | -- | -- | -- |
|  |  | Target | -- | -- | -- |
|  |  | Distractor | increase | -- | -- |
|  | Cz | Standard | increase | increase | OA > YA |
|  |  | Target | increase | increase | OA > YA |
|  |  | Distractor | increase | increase | OA > YA |
|  | Pz | Standard | increase | increase | -- |
|  |  | Target | increase | increase | -- |
|  |  | Distractor | increase | increase | -- |
| Resting Alpha Power | Fz |  | decrease | -- | OA > YA |
|  | Cz |  | decrease | -- | OA > YA |
|  | Pz |  | decrease | -- | -- |
| Aperiodic Exponent | Fz |  | decrease | -- | -- |
|  | Cz |  | decrease | -- | -- |
|  | Pz |  | decrease | increase | -- |
| Aperiodic Offset | Fz |  | decrease | increase | -- |
|  | Cz |  | decrease | increase | -- |
|  | Pz |  | decrease | increase | -- |

In the three-stimulus auditory oddball paradigm, the salient distractor stimulus elicits a P3a component (also known as “novelty P300”) that typically has a frontal maximum, while the infrequent target stimulus elicits a P3b component (also known as “target P300”) that has a centroparietal maximum (Polich, 2007). This distinct topographic distribution between the P3a and P3b component are based on different neural generative origins and activation patterns are a consequence of functionally different attention mechanisms (Polich, 2007). Interestingly, in this study, we observed age-related differences in the effects of arousal specifically on responses to the distractor tone, but not the target tone. The LC-NA system plays a central role in mediating multiple mechanisms of attentional control, and dysregulation of this neuromodulatory system has been associated with changes in these processes (Bouret & Sara, 2004, 2005; Sara & Bouret, 2012; Torres et al., 2025). Although both P3a and P3b components are modulated by LC-NA system activity, our findings indicate that only the P3a response to the distractor tone at frontal electrode sites was significantly modulated by arousal. This pattern reflects the common finding that elevated tonic LC activity is associated with heightened distractibility and more shifts in behavior (Bouret & Sara, 2005). The LC-NA system plays a critical role in modulating attentional priority by enhancing the salience of high-priority stimuli and suppressing low-priority stimuli through local hotspots produced by glutamatergic and noradrenergic feedback mechanisms (Mather et al., 2016). While we observed age-related differences in both components, the selective modulation of distractor responses under heightened arousal suggests a specific dysregulation of the LC-NA system that disproportionately affects rapid stimulus prioritization mechanisms rather than slower, goal-directed attentional processes that are linked to the facilitation of working memory. This interpretation aligns with neuroimaging findings indicating that individuals with elevated tonic arousal show greater disruptions in the frontoparietal attention networks, resulting in exaggerated processing of irrelevant stimuli through a model of adaptive gain (Aston-Jones & Cohen, 2005; Unsworth & Robison, 2017). Together, these findings suggest that LC-NA dysregulation in aging may bias attention dynamics toward heightened distractibility by shifting tonic LC activity to a suboptimal range for focused cognitive engagement.

Moreover, the inclusion of both passive and active task conditions in the present study highlights age-related differences in the underlying mechanisms of attentional control. Previous research has demonstrated that aging is associated with reduced P300 amplitudes to target stimuli and increased peak latencies in two-stimulus oddball paradigms (Mertens & Polich, 1997). Here, we employed a three-stimulus oddball design in which the distractor tone theoretically also elicits an earlier P3a component while the target tone primarily elicits only a P3b component (Polich, 2007). As expected, in our study, the P3a peak latencies elicited by the distractor tone in the frontal electrode was significantly faster than the P3b peak latencies elicited by the target tone in the central electrode (Friedman et al., 2001). In addition, our results reveal that older adults exhibit multiple signs of reduced inhibitory control. At frontal scalp sites, older adults showed robust P300 amplitudes across all stimulus types, whereas younger adults displayed attenuated responses. Although this frontal negativity in young adults may be a function of the centro-parietal distribution of the P300 dipole, the P300 component is also understood to represent inhibitory processing during stimulus discrimination, with a topographic shift from frontal to temporoparietal regions to facilitate working memory (Polich, 2007; Soltani & Knight, 2000). These processes are thought to be modulated by synaptic plasticity mechanisms involving both acetylcholine and noradrenaline (Ranganath & Rainer, 2003). Furthermore, the rapid habituation of the frontal P3a to novel stimuli has been interpreted as evidence of a suppression mechanism within stimulus evaluation systems (Friedman et al., 2001; Richardson et al., 2011). These mechanisms have been corroborated in recent magnetoencephalography study using a competitive attention task that identified reduced inhibition of irrelevant information and reduced allocation of cognitive resources that coincided with impaired distractor processing (ElShafei et al., 2022). Furthermore, schizophrenia research has consistently shown that reduced P3b amplitudes and abnormal P3a responses to distractors reflect impaired inhibitory control and attention filtering (Hamilton et al., 2024; Jiang et al., 2025; Mathalon et al., 2000). These patterns suggest that disruptions in stimulus discrimination, whether due to neurogenerative aging or psychiatric pathology, may share underlying mechanisms involving dysregulation of prefrontal networks and neuromodulatory systems. Our findings support the interpretation that while young adults exhibit optimal suppressive responses, older adults demonstrated impairments aligning with age-related declines in inhibitory function. In addition, at frontal, central, and parietal scalp locations, older adults displayed robust responses to frequent standard tones that were markedly weaker in young adults. Behaviorally, older adults did not exhibit slower response times to standard stimuli as would typically be predicted in the aging literature. This age equivalent effect likely reflects a diminished efficiency of frontal attentional control and increased attention allocation that is not evident in cognitively high-performing older adults (Daffner et al., 2007). These data support prior findings that older adults exhibit reduced habituation to consistently presented stimuli unlike young adults (ElShafei et al., 2022; Friedman et al., 1998; Getzmann et al., 2024). Although this lack of habituation and thus consequently increased distraction has primarily been identified in the frontal sites (Richardson et al., 2011), our findings reveal that this pattern extends throughout the scalp although most robust at the frontal electrode. Finally, older adults have been shown to exhibit longer P300 latencies in passive relative to active task conditions (Morgan & Murphy, 2010). These results align with our pupillometry findings which demonstrated that older adults exhibited enhanced pupil dilation responses to both target and distractor tones under active task conditions, as also seen in young adults, that mirror the faster P300 latencies observed during active attention processing in Morgan & Murphy (2010). Collectively, these findings suggest that increased distractibility, reduced habituation, and elevated attention allocation to salient but task-irrelevant stimuli in older adults may be driven by reduced frontal inhibitory control, leading to generalized overprocessing of all stimuli regardless of their attentional priority. Prior findings have shown that age-related mechanisms of filtering relevant from irrelevant information is a function of altered rhythmic neural activity in the 8-30 Hz oscillatory power range (Dahl, Ilg, et al., 2019). Future research should investigate whether these effects stem from delayed stimulus evaluation as seen in prolonged P300 latencies in older adults, which may be interacting with timely discrimination of stimuli and ultimately impairing the integration of inhibitory mechanisms.

Our findings also demonstrate that increased arousal differentially modulates resting-state EEG activity in young and older adults, revealing age-related dissociations across both oscillatory alpha dynamics and aperiodic spectral components. Consistent with prior studies, we observed an age-related slowing of individual peak alpha frequencies and reductions in relative alpha power across all scalp regions (Klimesch, 1999; Scally et al., 2018). Importantly, arousal selectively increased alpha power in older adults during the eyes-closed condition at frontal and central electrodes that was not observed in young adults. This suggests that alpha oscillations in the aging brain remain sensitive to state-level arousal but also reveal age-related changes in the role of the LC-NA system influence in these regions. The LC-NA system is known to regulate cortical network sensitivity through modulation of the attentional priority of inputs, specifically in the frontoparietal attention network through thalamocortical synchronization (Dahl, Mather, & Werkle-Bergner, 2022). Our data suggests that arousal more strongly impacts alpha dynamics in older adults, indicating potential compensatory or maladaptive desynchronization linked to altered LC-NA function. Prior studies have indeed shown that greater alpha desynchronization in aging is linked to cognitive decline (Dahl, Ilg, et al., 2019; Deiber et al., 2013; Leenders et al., 2018; Sander et al., 2012; Wöstmann et al., 2015). In addition, arousal differentially modulated aperiodic components extracted via spectral parameterization (Donoghue et al., 2020). In line with prior work, older adults in our study showed significantly flatter spectral slopes (i.e., lower aperiodic exponent), reflecting diminished inhibitory control (Donoghue et al., 2020; Merkin et al., 2023; Voytek et al., 2015). Furthermore, arousal similarly modulated both aperiodic exponent and offset in young and older adults. Increases in aperiodic exponent under arousal likely reflect a shift toward greater excitatory signaling, while elevations aperiodic offset may indicate a global shift in baseline neural firing or cortical activation.

In the context of neurodegenerative disease progression, individuals with mild cognitive impairment (MCI) have been shown to exhibit delayed P300 latencies and reduced amplitudes in response to targets tones compared with cognitive healthy older adults (Demirayak et al., 2023). Similar declines in the P300 component have also been consistently reported in a meta-analysis of patients with Alzheimer’s disease (Tarawneh et al., 2021). Interestingly, these latency delays are significantly more pronounced in the active version of oddball paradigms, suggesting that task-engaged conditions may enhance the sensitivity of the P300 to pathological cognitive decline (Knott et al., 1999). Given that our findings demonstrate a link between LC-NA system changes and P300 latencies, the P300 component may be able to serve as a sensitive biomarker for tracking progression from cognitively healthy stages to prodromal stages such as MCI and full-blown dementia. The field currently lacks a non-invasive, sensitive, and reliable functional measures to detect which asymptomatic or preclinical individuals are most at risk for progression toward dementia. Delayed intervention has been shown to be less effective in halting neurodegenerative process once they have significantly advanced (Yiannopoulou et al., 2019). For instance, in a clinical trial of Donanemab, an amyloid clearance antibody, patients with low to medium tau pathology showed greater than 20% slowing of cognitive decline on both the Integrated Alzheimer’s Disease Rating Scale and the Clinical Dementia Rating scale, compared with patients in the high tau pathology cohort (Sims et al., 2023). However, the greatest obstacle to successful prevention remains the early identification of asymptomatic adults who fall below threshold for current biomarkers. The present findings, in conjunction with prior research, suggest that behavioral, pupillometric, and scalp electrophysiological measures offer non-invasive and cost-effective tools for detection of early dysfunction in the LC-NA system in healthy aging, which may contribute to pathological spread.

Several limitations of the present study should be acknowledged. Due to methodological constraints in directly measuring LC neuronal firing in humans, we relied on indirect indices of LC-NA system activity including pupillometry and the P300 component of the event-related potential. Although both measures have been linked to noradrenergic function in humans (Gilzenrat et al., 2010; Murphy et al., 2011; Nieuwenhuis et al., 2005), they reflect the integrated output of multiple neural processes. Specifically, cholinergic activity also modulates pupil size. However, given the distinct temporal dynamics of these neuromodulatory systems, stimulus-evoked pupil responses are considered to more strongly index the rapid, phasic responses attributed to the LC-NA system (Reimer et al., 2016). While we employed many design elements to enhance the specificity of the LC-NA system (e.g., three-stimulus auditory oddball, threat of unpredictable shock), it remains unclear whether the observed effects reflect specific changes in neuronal firing or a summative effect of broader neuromodulatory dynamics. In particular, orexin neurons in the lateral hypothalamic area also play a key role in arousal regulation and degenerate early, but their specific contributions to our findings remain unclear (Oh et al., 2019). Another methodological consideration is comparison of P300 components between age groups. We *a priori* focused on the Fz, Cz, and Pz electrodes given prior evidence of maximal P300 amplitudes along the midline (Mertens & Polich, 1997; Polich, 1987, 2007). However, age-related changes in scalp topography may warrant alternative approaches such as cluster-based permutation testing or individualized peak selection, particularly in the presence of hemispheric asymmetries, which has also revealed similar age-related patterns to target and distractor tone processing (Getzmann et al., 2024). In addition, our use of a fixed white noise burst as a distractor stimulus may have limited our ability to capture novelty-related LC-NA responses and may better be associated with saliency-related LC-NA responses. Future studies may benefit from using unique distractor stimuli on each trial to better engage phasic LC activity, although similar age-related impairments have previously been observed (ElShafei et al., 2022).

Finally, we interpret our findings based on rodent experiments demonstrating a negative inverse relationship between LC tonic and phasic responses (Hayat et al., 2020). Our study employed indirect indices of phasic noradrenergic responses (i.e., P300, pupillometry), under the assumption that age-related changes in tonic activity would be reflected in these measures. However, we note two key considerations. First, the linear relationship between tonic and phasic activity has not been proven in humans and older adults may exhibit altered phasic responses independent of tonic baseline levels. For instance, baseline pupil size, an index of tonic activity, shows a monotonic decline with age, but phasic pupil responses follow an inverted-U pattern across the lifespan (Riley et al., 2024). This dissociation suggests that aging may not preserve this strict linear relationship and that aging may uniquely alter the dynamics of both tonic activity and phasic responses. Second, as previously mentioned, these indirect measures are not specific to noradrenergic function, raising questions about their sensitivity and reliability to capture perfect tonic-phasic relationships. Given these considerations, we adopted this unique multimodal approach and manipulated arousal within subjects to test our hypotheses. Our findings suggest that behavioral and electrophysiogical measures may be more sensitive to investigate changes in noradrenergic function compared with pupillometry, oscillatory signals, and aperiodic activity. This study further emphasizes the need for absolute and reliable measures of LC function in humans to assess important questions in aging and neurodegenerative diseases.

The present study provides multimodal evidence that arousal regulation and attentional control differ between cognitively healthy younger and older adults. We observed consistent patterns indicating that older adults are more susceptible to increased distractibility and diminished stimulus discrimination, particularly under heightened arousal. These findings support theoretical models that age-related changes in tonic LC activity impair phasic responsiveness, leading to reduced behavioral flexibility in modulating attention and cortical excitability. Interestingly, we observed age-equivalent modulation of aperiodic spectral features by arousal that suggests some preservation of cortical neuromodulatory responsiveness to arousal, even as older adults displayed significant alterations in oscillatory dynamics. Our results also highlight the value of passive and active task conditions in revealing latent age-related differences in attentional engagement and inhibitory control. In addition to our electrophysiology results, the most striking findings emerged from the pupillometry data. During passive listening, older adults exhibited pupil responses comparable to those of young adults. However, during active task engagement, significant age-related differences appeared for the standard and distractor tones, but not for the target tone. This pattern indicates that age differences in phasic noradrenergic activity depend on multiple contextual factors including state (resting vs. task engaged), arousal (neutral vs. heightened), and the type of attention processing (suppression of task-irrelevant stimuli, selective attention to goal-directed stimuli, and attention capture by salient stimuli). These results may reflect altered LC-NA system phasic responsivity across conditions in older adults. Finally, we replicated age-related changes in P300 latencies, alpha dynamics, and aperiodic signals that may serve as sensitive, non-invasive biomarkers of early neuromodulatory dysfunction. These functional markers hold potential for identifying individuals at elevated risk for neurodegenerative decline, prior to the presence of existing biomarkers of neurodegeneration at preclinical stages of AD. Future longitudinal research should further evaluate these indices to track trajectories of cognitive aging and determine their predictive value in early detection, that may lead to interventions aimed to perverse LC-NA system function in late life and impede the progression of neurodegenerative diseases.

## Author Contributions

AJK and MM designed the experiment. AJK and JS collected all data. AJK and SM created the analysis code. AJK conducted all data analyses. JS contributed to data curation. AJK wrote the initial manuscript. All coauthors revised the manuscript and approved the final version.

