## Supplemental Table 1 for "Electroencephalography, pupillometry, and behavioral evidence for locus coeruleus-noradrenaline system related tonic hyperactivity in older adults"

|  | **Condition** | **Standard** | **Target** | **Distractor** |
| --- | --- | --- | --- | --- |
| **Passive Task** | **Young Adults** | | | |
|  | No-threat | 6.02 (0.48) | 6.13 (0.48) | 7.14 (0.65) |
|  | Threat | 6.66 (0.52) | 8.65 (2.31) | 8.11 (0.86) |
|  | **Older Adults** | | | |
|  | No-threat | 4.74 (0.59) | 4.33 (0.59) | 5.74 (0.86) |
|  | Threat | 5.05 (0.55) | 6.46 (1.20) | 6.90 (1.68) |
| **Active Task** | **Young Adults** | | | |
|  | No-threat | 5.74 (0.43) | 8.09 (0.67) | 6.83 (0.54) |
|  | Threat | 6.54 (0.47) | 7.69 (0.60) | 7.61 (0.65) |
|  | **Older Adults** | | | |
|  | No-threat | 3.60 (0.37) | 5.96 (0.81) | 5.31 (0.74) |
|  | Threat | 4.15 (0.43) | 6.82 (0.87) | 5.63 (0.66) |
