## Supplemental Table 2 for "Electroencephalography, pupillometry, and behavioral evidence for locus coeruleus-noradrenaline system related tonic hyperactivity in older adults"

|  | **Condition** | **Standard** | **Target** | **Distractor** |
| --- | --- | --- | --- | --- |
| *Fz Electrode* | | | | |
| **Passive Task** | **Young Adults** | | | |
|  | No-threat | -0.50 (0.22) | -1.63 (0.33) | -1.12 (0.42) |
|  | Threat | -0.17 (0.21) | -1.35 (0.33) | -0.21 (0.24) |
|  | **Older Adults** | | | |
|  | No-threat | -0.44 (0.13) | 0.03 (0.27) | 0.45 (0.32) |
|  | Threat | -0.29 (0.16) | -0.13 (0.29) | 0.20 (0.32) |
| **Active Task** | **Young Adults** | | | |
|  | No-threat | 0.14 (0.61) | -1.05 (0.57) | -0.29 (0.57) |
|  | Threat | -0.21 (0.65) | -2.16 (0.56) | -0.69 (0.47) |
|  | **Older Adults** | | | |
|  | No-threat | 3.03 (0.51) | 2.21 (0.54) | 2.04 (0.51) |
|  | Threat | 2.42 (0.50) | 1.66 (0.54) | 1.68 (0.46) |
| *Cz Electrode* | | | | |
| **Passive Task** | **Young Adults** | | | |
|  | No-threat | -0.29 (0.13) | -0.74 (0.21) | 0.40 (0.36) |
|  | Threat | -0.12 (0.16) | -0.98 (0.23) | 0.81 (0.24) |
|  | **Older Adults** | | | |
|  | No-threat | -0.45 (0.12) | 0.25 (0.29) | 0.91 (0.35) |
|  | Threat | -0.25 (0.12) | 0.01 (0.24) | 0.83 (0.30) |
| **Active Task** | **Young Adults** | | | |
|  | No-threat | 1.03 (0.37) | 3.13 (0.48) | 2.85 (0.43) |
|  | Threat | 0.38 (0.37) | 1.94 (0.44) | 2.17 (0.40) |
|  | **Older Adults** | | | |
|  | No-threat | 3.44 (0.38) | 3.50 (0.53) | 2.69 (0.46) |
|  | Threat | 2.79 (0.41) | 2.79 (0.56) | 2.31 (0.47) |
| *Pz Electrode* |  |  |  |  |
| **Passive Task** | **Young Adults** | | | |
|  | No-threat | 0.02 (0.12) | 0.09 (0.21) | 1.01 (0.26) |
|  | Threat | -0.32 (0.12) | -0.55 (0.22) | 1.07 (0.30) |
|  | **Older Adults** | | | |
|  | No-threat | -0.34 (0.09) | 0.10 (0.26) | 0.69 (0.30) |
|  | Threat | -0.27 (0.09) | 0.14 (0.21) | 0.72 (0.28) |
| **Active Task** | **Young Adults** | | | |
|  | No-threat | 0.64 (0.29) | 4.64 (0.49) | 3.66 (0.48) |
|  | Threat | 0.53 (0.32) | 4.23 (0.46) | 3.22 (0.42) |
|  | **Older Adults** | | | |
|  | No-threat | 1.88 (0.36) | 3.01 (0.49) | 2.27 (0.41) |
|  | Threat | 1.91 (0.32) | 3.02 (0.49) | 2.32 (0.42) |
