## Supplemental Table 3 for "Electroencephalography, pupillometry, and behavioral evidence for locus coeruleus-noradrenaline system related tonic hyperactivity in older adults"

|  | **Condition** | **Standard** | **Target** | **Distractor** |
| --- | --- | --- | --- | --- |
| *Fz Electrode* | | | | |
| **Passive Task** | **Young Adults** | | | |
|  | No-threat | 390.19 (7.84) | 385.28 (7.81) | 362.54 (7.95) |
|  | Threat | 388.80 (6.19) | 384.81 (7.39) | 364.67 (7.41) |
|  | **Older Adults** | | | |
|  | No-threat | 374.04 (7.53) | 385.19 (6.91) | 380.74 (6.00) |
|  | Threat | 378.24 (7.08) | 391.48 (7.20) | 387.96 (7.59) |
| **Active Task** | **Young Adults** | | | |
|  | No-threat | 362.94 (5.93) | 355.61 (6.96) | 348.51 (6.50) |
|  | Threat | 368.80 (5.31) | 354.98 (5.72) | 345.20 (5.89) |
|  | **Older Adults** | | | |
|  | No-threat | 372.81 (5.78) | 369.35 (5.71) | 375.56 (6.91) |
|  | Threat | 380.54 (6.16) | 383.57 (5.89) | 391.39 (7.69) |
| *Cz Electrode* | | | | |
| **Passive Task** | **Young Adults** | | | |
|  | No-threat | 400.65 (6.13) | 380.37 (8.71) | 375.74 (8.27) |
|  | Threat | 415.37 (7.01) | 406.02 (8.47) | 389.91 (8.55) |
|  | **Older Adults** | | | |
|  | No-threat | 404.54 (6.69) | 399.91 (6.10) | 399.35 (6.65) |
|  | Threat | 395.19 (6.95) | 395.56 (6.97) | 401.48 (7.27) |
| **Active Task** | **Young Adults** | | | |
|  | No-threat | 382.69 (7.46) | 372.50 (8.17) | 369.72 (8.80) |
|  | Threat | 397.87 (6.88) | 382.87 (7.77) | 373.06 (9.43) |
|  | **Older Adults** | | | |
|  | No-threat | 395.65 (6.16) | 388.11 (6.00) | 386.02 (6.61) |
|  | Threat | 422.13 (5.38) | 407.22 (6.51) | 413.98 (7.99) |
| *Pz Electrode* |  |  |  |  |
| **Passive Task** | **Young Adults** | | | |
|  | No-threat | 390.76 (6.97) | 388.98 (7.79) | 379.63 (7.73) |
|  | Threat | 399.07 (7.99) | 397.83 (7.45) | 396.67 (8.83) |
|  | **Older Adults** | | | |
|  | No-threat | 403.61 (6.48) | 406.39 (6.73) | 402.87 (6.47) |
|  | Threat | 410.65 (6.91) | 412.78 (6.68) | 410.00 (6.57) |
| **Active Task** | **Young Adults** | | | |
|  | No-threat | 382.31 (8.04) | 393.98 (8.15) | 381.76 (8.70) |
|  | Threat | 387.50 (7.36) | 392.13 (7.04) | 387.41 (7.80) |
|  | **Older Adults** | | | |
|  | No-threat | 411.17 (6.12) | 403.33 (6.66) | 411.76 (7.53) |
|  | Threat | 420.28 (6.07) | 421.20 (6.34) | 426.02 (6.91) |
