## Supplemental Table 4 for "Electroencephalography, pupillometry, and behavioral evidence for locus coeruleus-noradrenaline system related tonic hyperactivity in older adults"

|  | **Condition** | **Alpha Power** | **Alpha Frequency** | **Aperiodic Exponent** | **Aperiodic Offset** |
| --- | --- | --- | --- | --- | --- |
| *Fz Electrode* | | | | |  |
| **Eyes Closed** | **Young Adults** | | | |  |
|  | No-threat | 87.9 (0.95) | 10.3 (0.12) | 1.54 (0.04) | 1.32 (0.05) |
|  | Threat | 87.0 (1.00) | 10.3 (0.12) | 1.58 (0.04) | 1.39 (0.05) |
|  | **Older Adults** | | | |  |
|  | No-threat | 79.8 (1.33) | 9.80 (0.14) | 1.38 (0.04) | 1.15 (0.06) |
|  | Threat | 80.8 (1.33) | 9.61 (0.12) | 1.42 (0.04) | 1.27 (0.06) |
| **Eyes Open** | **Young Adults** | | | |  |
|  | No-threat | 79.1 (1.21) | 10.5 (0.13) | 1.44 (0.04) | 1.19 (0.06) |
|  | Threat | 82.3 (1.17) | 10.3 (0.12) | 1.42 (0.04) | 1.22 (0.06) |
|  | **Older Adults** | | | |  |
|  | No-threat | 71.1 (1.54) | 9.67 (0.16) | 1.26 (0.05) | 1.00 (0.07) |
|  | Threat | 72.9 (1.60) | 9.45 (0.14) | 1.24 (0.05) | 1.08 (0.06) |
| *Cz Electrode* | | | | |  |
| **Eyes Closed** | **Young Adults** | | | |  |
|  | No-threat | 87.6 (0.96) | 10.2 (0.11) | 1.62 (0.04) | 1.12 (0.08) |
|  | Threat | 87.4 (0.97) | 10.3 (0.11) | 1.67 (0.04) | 1.23 (0.07) |
|  | **Older Adults** | | | |  |
|  | No-threat | 80.3 (1.34) | 9.70 (0.13) | 1.41 (0.04) | 0.99 (0.07) |
|  | Threat | 81.5 (1.30) | 9.66 (0.12) | 1.46 (0.04) | 1.14 (0.07) |
| **Eyes Open** | **Young Adults** | | | |  |
|  | No-threat | 79.6 (1.20) | 10.3 (0.13) | 1.47 (0.04) | 0.90 (0.07) |
|  | Threat | 83.2 (1.14) | 10.2 (0.13) | 1.52 (0.04) | 1.05 (0.07) |
|  | **Older Adults** | | | |  |
|  | No-threat | 71.7 (1.57) | 9.90 (0.17) | 1.25 (0.05) | 0.78 (0.07) |
|  | Threat | 74.4 (1.60) | 9.54 (0.12) | 1.27 (0.04) | 0.91 (0.06) |
| *Pz Electrode* |  |  |  |  |  |
| **Eyes Closed** | **Young Adults** | | | |  |
|  | No-threat | 88.5 (0.86) | 10.5 (0.13) | 1.57 (0.04) | 0.97 (0.07) |
|  | Threat | 89.0 (0.82) | 10.5 (0.12) | 1.61 (0.04) | 1.02 (0.07) |
|  | **Older Adults** | | | |  |
|  | No-threat | 82.3 (1.29) | 10.1 (0.13) | 1.30 (0.04) | 0.68 (0.07) |
|  | Threat | 83.2 (1.27) | 9.95 (0.12) | 1.37 (0.04) | 0.88 (0.07) |
| **Eyes Open** | **Young Adults** | | | |  |
|  | No-threat | 82.4 (1.16) | 10.5 (0.13) | 1.46 (0.04) | 0.78 (0.07) |
|  | Threat | 86.9 (1.03) | 10.3 (0.12) | 1.50 (0.04) | 0.91 (0.06) |
|  | **Older Adults** | | | |  |
|  | No-threat | 73.9 (1.64) | 10.2 (0.15) | 1.16 (0.05) | 0.48 (0.08) |
|  | Threat | 77.6 (1.65) | 9.84 (0.11) | 1.21 (0.05) | 0.64 (0.07) |
